# PfApiAT2 is a proline transporter essential for the transmission of *Plasmodium falciparum* by the mosquito vector

**DOI:** 10.64898/2026.01.20.700404

**Authors:** Malhar Khushu, R. Charles Kissel, Jamie Kauffman, Claudia Taccheri, Naresh Singh, Robert L. Summers, Leigh D. Plant, Dyann F. Wirth, Flaminia Catteruccia, Selina Bopp

## Abstract

*Plasmodium falciparum* oocysts undergo an explosive biomass increase during development in *Anopheles* mosquitoes, a dramatic growth process likely promoted by as-yet unknown nutrients scavenged from the mosquito. We previously observed in blood-stage parasites, that the amino acid transporter PfApiAT2, although dispensable, regulates proline homeostasis and mediates resistance to halofuginone, a potent proline-tRNA synthetase inhibitor. Here, we demonstrate that PfApiAT2 is a proline-specific transporter essential for early oocyst development in *Anopheles gambiae*. Halofuginone-resistant *pfapiat2*-mutant parasites form stunted oocysts severely defective in sporozoite production. This phenotype is recapitulated in PfApiAT2-knockout parasites that undergo a complete block in sporogony, forming oocysts that stall and degenerate. Remarkably, this growth defect can be rescued by nutrient supplementation to the mosquito vector. By identifying an amino acid transporter essential for oocyst growth, our data unveil a vulnerability in *P. falciparum* transmission, revealing a critical nutritional dependency of the parasite on its mosquito vector.

## Introduction

Malaria remains a global public health crisis, causing 263 million cases in 2024 [1]. The *Plasmodium* parasites that cause this disease have an intricate lifecycle spanning a multitude of tissue environments, performing complex morphological changes and massive biomass expansion in both vertebrate and invertebrate hosts. These parasites are dependent on the human host and insect vector to scavenge nutrients like amino acids for growth. During the intraerythrocytic asexual stage in the human host, parasites obtain most of their amino acids by breakdown of hemoglobin derived from the host red blood cell they have invaded, and a minority through transporter-mediated uptake from the host plasma [2]. Nutrient acquisition during other stages of the parasite’s lifecycle is poorly understood, especially during development in the mosquito vector, where oocysts grow over a period of 7-10 days while embedded under the midgut basal lamina. Here, parasites are bathed in the hemolymph, the mosquito open circulatory system, from which they are thought to scavenge nutrients to facilitate their explosive expansion and subsequent division into many thousands of daughter sporozoites [3, 4]. The reliance of oocysts on mosquito nutrient stores and their ability to scavenge nutrients from the mosquito environment under changing nutritional conditions has been under much investigation [5–12]. Despite progress, however, the importance of free amino acid acquisition during this period of growth has been largely unexplored [13].

The genome of *Plasmodium falciparum*, one of the major causative agents of malaria, encodes a relatively short list of membrane transporters (only 2.1% of *P. falciparum* genes, compared with 4.2% for *Saccharomyces cerevisiae* and 3.4% for *Homo sapiens*) [14]. Of these, some peptide and amino acid transporters (for eg. *Pf*CRT (Chloroquine Resistance Transporter) and *Pf*AAT1 (Amino Acid Transporter 1)) have been associated with mutations conferring resistance to frontline antimalarials [15, 16]. The function of these and other transporters has primarily been described during the asexual blood-stages, with limited work on the function of parasite amino acid transporters in the mosquito [17].

We have recently discovered that a putative amino acid transporter, PfApiAT2 (Apicomplexan Amino acid Transporter 2, also known as MFR4, Major Facilitator superfamily protein 4) is involved in resistance to halofuginone (HFG), a potent proline-competitive inhibitor of the parasite cytoplasmic proline-tRNA synthase (cPRS). Parasites lacking PfApiAT2 function (via either selection of loss-of-function mutants or genetic knockouts) show no observable defects during the asexual blood-stage, but have vastly increased proline levels in the parasite cytoplasm (a mechanism initially described as the Adaptive Proline Response, APR) and thus are thought to outcompete HFG for binding at the target site [18–21]. PfApiAT2 is part of the ApiAT family of Apicomplexan Amino acid Transporters, initially described as Novel Putative Transporters (NPTs)[14]. ApiATs belong to the Major Facilitator superfamily of proteins, with 12 transmembrane domains, and canonical polytopic membrane-transversing folds for performing facilitated diffusion down a concentration gradient. The substrates of several ApiATs have been discovered; in *Toxoplasma* gondii, TgApiAT5-3 is described as a transporter of large aromatic amino acids, primarily tyrosine, and TgApiAT1 (TgNPT1) is reported to mediate arginine and lysine transport [22, 23]. In multiple *Plasmodium* species, ApiAT8 (NPT1) is reported to transport arginine, lysine and other cationic metabolites, and has been associated with essential functions during establishment of infection in the mosquito vector [22, 24, 25]. Moreover, in the mouse malaria parasite *Plasmodium berghei*, PbApiAT2 is essential for proper sporogonic development in the mosquito by unknown mechanisms, and its substrate/s is also unknown [17].

Here we demonstrate that HFG-resistant (HFG-R) *P. falciparum* parasites with non-synonymous mutations in the *pfapiat2* locus exhibit a severe fitness cost to transmission through female *Anopheles gambiae* mosquitoes, a major vector in sub-Saharan Africa. Additionally, we find that *P. falciparum* parasites with a genetically ablated *pfapiat2* locus (PfApiAT2-KO) are similarly unable to complete sporogony. Using a combination of immunofluorescence and transmission electron microscopy, we determine that PfApiAT2 is expressed on the plasma membrane of early oocysts, and that its genetic knockout stalls oocyst growth eventually leading to their degeneration. With a *Xenopus laevis* oocyte heterologous expression system, we show that PfApiAT2 is a proline-specific transporter. Finally, we determine that PfApiAT2-KO parasites can be rescued via supplementation of exogenous amino acids and an additional blood meal to the mosquito vector. These findings demonstrate that proline transport by PfApiAT2 is essential for oocyst growth in the mosquito vector, revealing a potential new target for transmission-blocking strategies.

## Results

### Halofuginone-resistant *pfapiat2* mutants are stunted in oocyst growth

Our past work in non-transmissible Dd2 and 3D7 parasite lines implicated PfApiAT2 loss-of-function in HFG-R parasites [21]. Previous work in *P. berghei* suggested that PbApiAT2 may play a role in mosquito stage transmission [17]. To investigate the fitness-cost of HFG resistance during transmission, resistant parasites with mutations in the *pfapiat2* locus were selected by culturing asexual blood-stages of NF54 (a *P. falciparum* strain readily transmissible by *Anopheles* females) under HFG pressure as previously described [21]. We recovered two bulk asexual stage cultures with different single-nucleotide polymorphisms (R345I and G449R) in the *pfapiat2* locus. These mutations, although distinct from previously reported SNPs selected under HFG selections [21], showed the characteristic ∼10-fold shift in HFG EC_50_ compared to wildtype (WT), reminiscent of parasites exhibiting APR, or PfApiAT2 genetic knockout parasites (Supplementary Figure 1). This indicated that R345I and G449R mutations likely cause a complete loss-of-function of PfApiAT2 [21]. We next determined whether these mutants are capable of development during mosquito stages. We induced WT and both mutant cultures to produce gametocytes, and used these gametocyte cultures to infect female *An. gambiae* mosquitoes (Figure 1A). At 7 days post-infection (dpi), both HFG-R parasite lines showed comparable infection of the mosquito midgut with similar oocyst numbers and prevalence compared to WT, although we observed a small but significant reduction in prevalence of G449R oocysts (Figure 1B, oocysts/midgut WT vs R345I *p*=0.0918, WT vs G449R *p*>0.9999, Kruskal-Wallis; percent infected mosquito midguts WT vs R345I *p*=0.2209, WT vs G449R *p*=0.0089, two-tailed Fisher’s Exact). However, we observed a striking reduction in oocyst size (Figure 1C), with average mutant oocyst area per midgut being 4-4.5 times smaller than WT for both mutant lines (WT vs R345I *p*<0.0001, WT vs G449R *p*< *p*<0.0001, Kruskal-Wallis). Growth over an additional 7 or 14 days did not result in an increase in the size of the overall population of mutant oocysts, indicating that parasites were either dead or permanently stalled (Supplementary Figure 2). Consistent with this finding, at 14dpi, the burden of salivary gland sporozoites in mosquitoes infected with either mutant line was far lower than in controls (percent infected mosquitoes 14dpi WT vs R345I *p*<0.0001, WT vs G449R *p*<0.0001, two-tailed Fisher’s Exact) (Figure 1D, left). Sporozoite prevalence and intensity remained low after an additional week of development (percent infected mosquitoes 21dpi WT vs R345I *p*<0.0001, WT vs G449R *p*<0.0001, two-tailed Fisher’s Exact; sporozoites/mosquito WT vs R345I *p*<0.0001, WT vs G449R *p*<0.0001, Kruskal-Wallis), although we observed a marginal increase in sporozoite prevalence in mosquitoes infected with G449R parasites relative to 14dpi (G449R 14dpi vs 21dpi *p*=0.0287, two-tailed Fisher’s Exact). These data suggest that mutations in the *pfapiat2* locus confer a fitness cost to parasites during oocyst growth and hint at an essential function for PfApiAT2 during this lifecycle stage.

**Figure 1:**
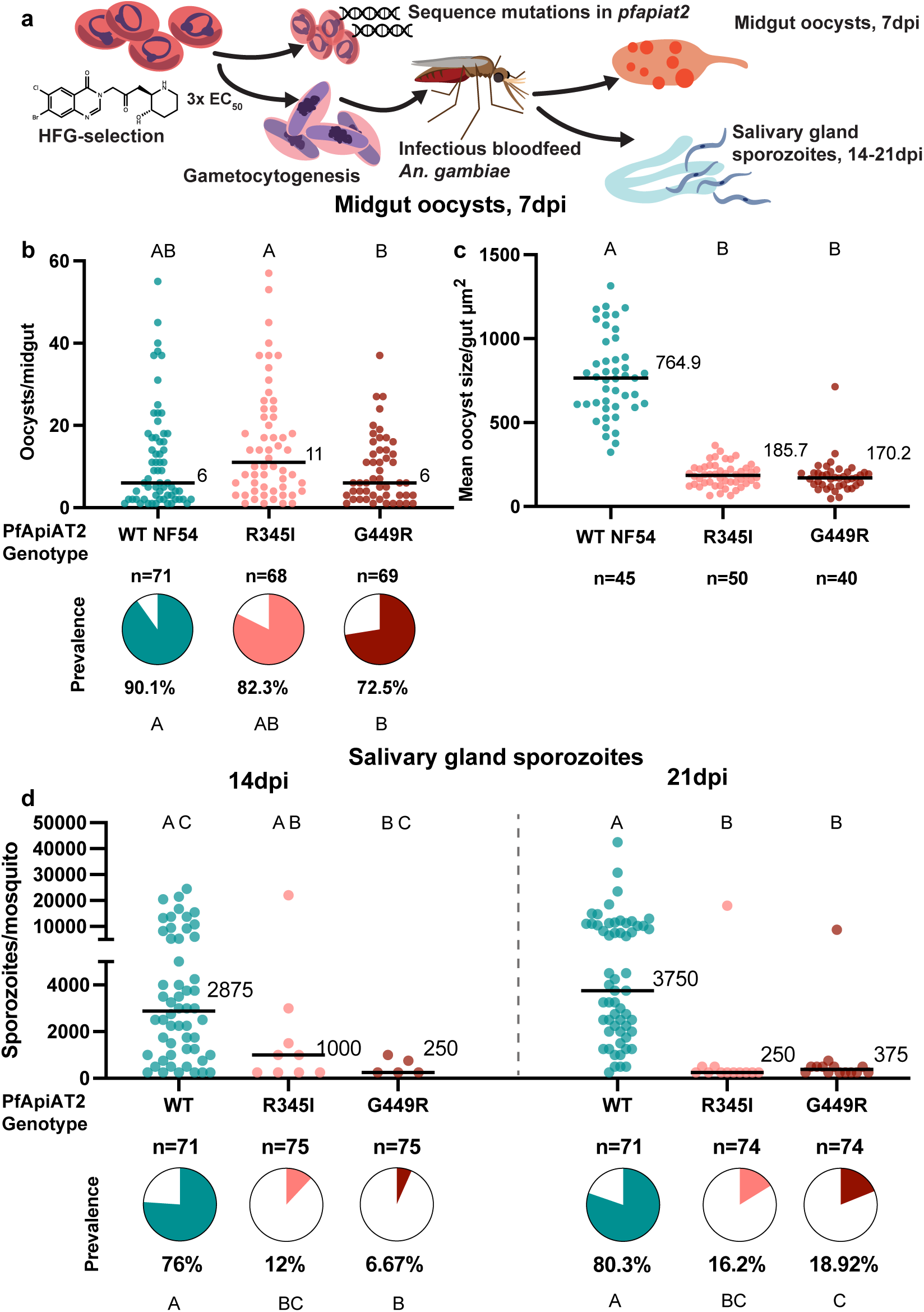
Halofuginone resistant parasites with mutations in the *pfapiat2* locus have defective oocyst development and fewer sporozoite numbers. (a) Scheme of selection of resistant parasites, followed by confirmation of mutations in *pfapiat2* via Sanger sequencing and transmission to female *An. gambiae* mosquitoes. (b) HFG-R parasites infect the mosquito midgut comparable to age-matched parental WT NF54 parasites (oocysts/midgut WT vs R345I *p*=0.0918; WT vs G449R *p*>0.9999, Kruskal-Wallis with post-hoc Dunn’s correction; % infected midguts WT vs R345I *p*=0.2209, WT vs G449R *p*=0.0089, two-tailed Fisher’s Exact test). Each dot represents number of oocysts per midgut, and line represents median. Pie charts represent % midguts with >1 oocyst. Data is pooled from 3 independent biological replicates. (c) HFG-R parasites form oocysts significantly smaller than WT NF54 parasites (WT vs R345I *p*<0.0001; WT vs G449R *p*<0.0001, Kruskal-Wallis with post-hoc Dunn’s correction). Each dot represents average area of oocysts per midgut from midguts with > 2 midguts, and line represents median. Data is pooled from 3 independent biological replicates. (d). Dissections of salivary glands at 14dpi and 21dpi revealed that salivary gland sporozoite burden of HFG-R parasites are starkly reduced at 14dpi (left) and 21dpi (right) (% infected mosquitoes 14dpi: WT vs R345I *p*<0.0001, WT vs G449R *p*<0.0001; 21dpi: WT vs R345I *p*<0.0001, WT vs G449R *p*<0.0001, two-tailed Fisher’s Exact test). A marginal increase in prevalence of sporozoites in mosquitoes infected with G449R-parasites between 14dpi and 21dpi is observed (% infected mosquitoes G449R 14dpi vs 21dpi *p*=0.0287, two-tailed Fisher’s Exact test). Each dot represents sporozoites counted via hemocytometer per dissected mosquito, and line represents median. Data is pooled from 3 independent biological replicates. Different letters indicate statistically significant differences, details of statistical tests and all comparisons tested provided in Supplementary Data. Source data are provided as a Source Data file.

### Genetic ablation of *pfapiat2* results in a complete block in sporogony

To directly confirm that the observed effects on parasite development were due to mutations in *pfapiat2*, we carried out a genetic knockout (KO) of the locus, using homology directed repair via CRISPR-Cas9 as previously done in non-transmissible Dd2 parasites [21]. The integration of the selectable marker was confirmed via PCR amplification around integration sites (Supplementary Figure 3A). As expected, PfApiAT2-KO parasites showed a shift in EC_50_ under drug pressure in asexual stages compared to WT (Supplementary Figure 3B). We then fed WT and PfApiAT2-KO gametocytes to *An. gambiae* females (Figure 2A), and dissected midguts and salivary glands over the duration of the infection. As observed with the HFG-R mutants, PfApiAT2-KO parasites established infection in the mosquito midgut (Figure 2B). However, the oocysts formed were 5 times smaller than WT parasites (Figure 2C, WT vs PfApiAT2-KO *p*<0.0001, Mann-Whitney), and showed an almost complete lack of sporozoites in the mosquito salivary glands at 14dpi (Figure 2D (left), percent infected mosquitoes 14dpi WT vs PfApiAT2-KO *p*<0.0001, two-tailed Fisher’s Exact), closely resembling the phenotypes observed in HFG-R parasites. Growth over an additional 7 days, until 21dpi, did not increase the salivary gland sporozoite burden (Figure 2D (right), percent infected mosquitoes 21dpi WT vs PfApiAT2-KO *p*<0.0001, two-tailed Fisher’s Exact). Combined, these data demonstrate that *pfapiat2*, a dispensable gene during asexual stages and gametocytogenesis, is a key transporter for completion of sporogony.

**Figure 2:**
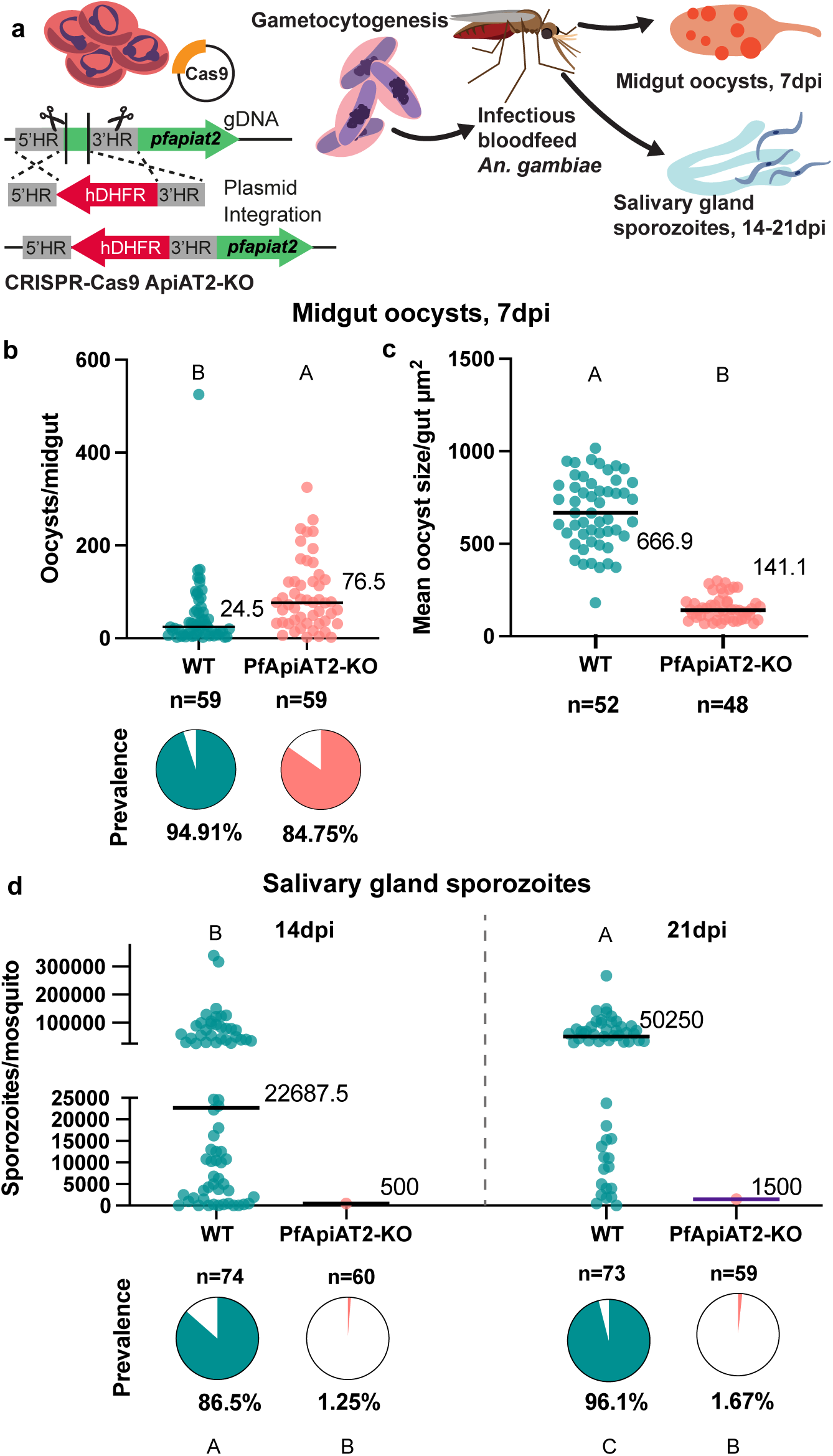
PfApiAT2-KO parasites form stunted oocysts that are unable to undergo sporogony. (a) Scheme of genetic disruption of PfApiAT2 using CRISPR-Cas9 by insertion of an hDHFR cassette into the *pfapiat2* locus in asexual stage NF54 parasites, followed by induction to mature, transmissible gametocytes and feeding to female *An. gambiae* mosquitoes. (b, c) (b) PfApiAT2-KO oocysts infect the midgut of *An. gambiae* females better than parental WT (oocysts/mosquito *p*<0.0001, Mann-Whitney), but (c) are almost 5 times smaller in size at day 7 post infection (*p*<0.0001, Mann-Whitney). (b) Each dot represents number of oocysts per midgut, and line represents median. Pie charts represent % midguts with >1 oocyst. Data is pooled from 4 independent biological replicates. (c) Each dot represents average area of oocysts per midgut from midguts with > 2 midguts, and line represents median. Data is pooled from 4 independent biological replicates. (d) Dissections of salivary glands at 14dpi (left) and 21dpi (right) revealed an almost complete lack of sporozoites in mosquitoes fed PfApiAT2-KO parasites, recapitulating the fitness cost observed in HFG-R parasites (% infected mosquitoes WT vs PfApiAT2-KO 14dpi P<0.0001; 21dpi *p*<0.0001, two-tailed Fisher’s Exact test). Each dot represents sporozoites counted via hemocytometer per dissected mosquito, and line represents median. Data is pooled from 3-4 independent biological replicates. Different letters indicate statistically significant differences, details of statistical tests and all comparisons tested provided in Supplementary Data. Source data are provided as a Source Data file.

### PfApiAT2-KO oocysts arrest during early growth

To pinpoint the dynamics of growth arrest, we measured the size of WT and PfApiAT2-KO oocysts between 3 to 6dpi, covering a considerable period of oocyst growth. We observed that PfApiAT2-KO oocysts deviate from WT in size starting at 4dpi (Figure 3A, mean oocyst size/midgut 3dpi WT vs PfApiAT2-KO *p*=0.6517, 4dpi WT vs PfApiAT2-KO *p*<0.0001, Kruskal-Wallis). While WT parasites continued in their exponential growth from 4dpi to 6dpi, we observed that PfApiAT2-KO oocyst growth began to plateau at 4dpi. We next determined whether this timing corresponded with expression of PfApiAT2, using transgenic NF54 parasites expressing PfApiAT2 tagged with the high-avidity spaghetti monster hemagglutinin (smHA) epitope tag, generated using CRISPR-Cas9 [26, 27] (Supplementary Figure 4A, B).

**Figure 3:**
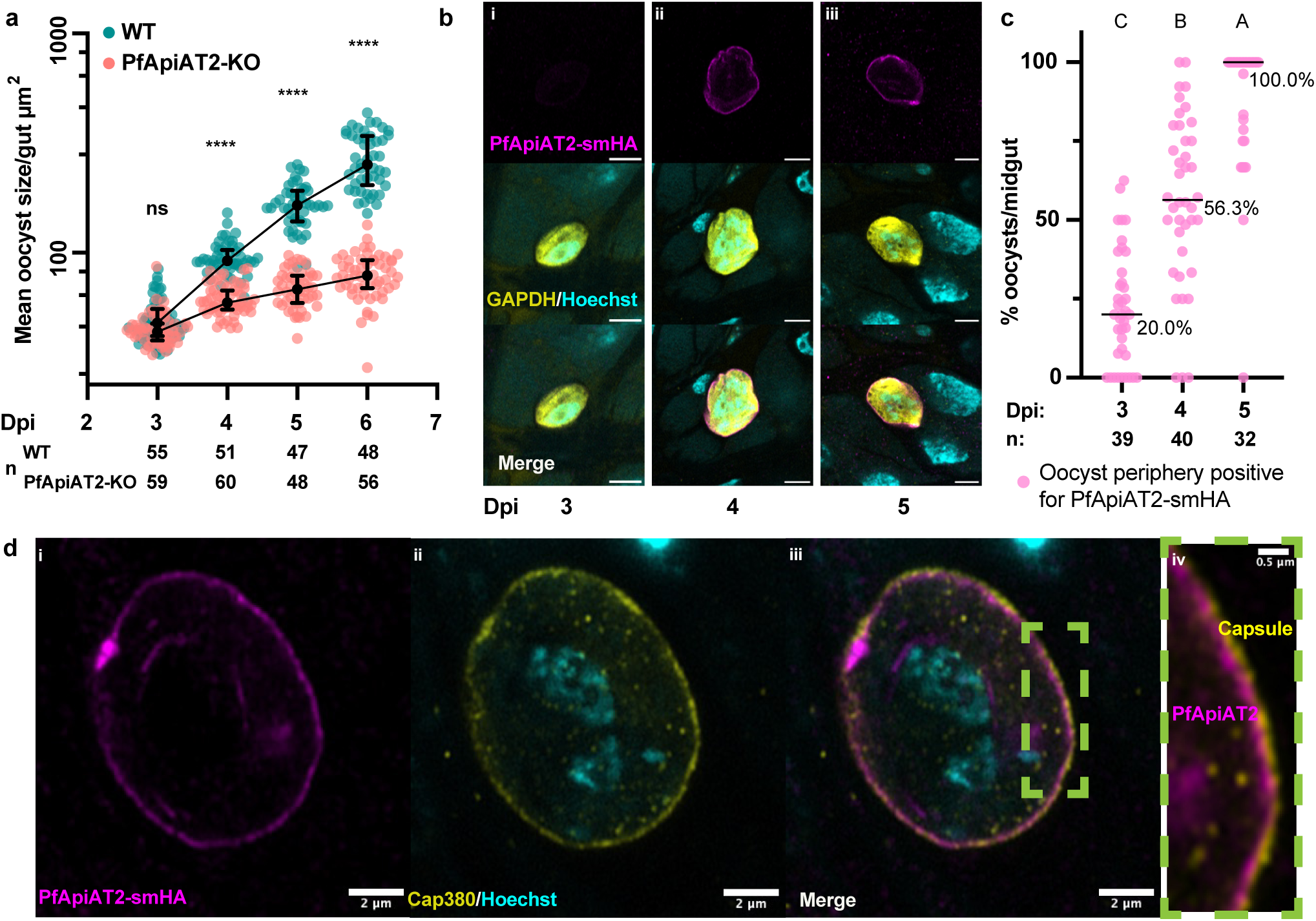
PfApiAT2-KO oocyst growth defect times closely with PfApiAT2 expression at the parasite plasma membrane. (a) WT and PfApiAT2-KO oocyst size from 3 to 6dpi. PfApiAT2-KO oocysts deviate in size from WT at 4dpi (ns: not significant, ****: *p*<0.0001, Kruskal-Wallis with post-hoc Dunn’s correction). Each dot represents average area of oocysts per midgut from midguts with > 2 midguts, and line represents median. Data is pooled from 3 independent biological replicates. (b) representative IFAs of PfApiAT2-smHA oocysts showing no smHA-signal at the parasite periphery at 3dpi, but present from 4dpi onwards. Scale bars are each 5μm (c) Quantification of PfApiAT2-smHA positivity at the parasite periphery revealed increasing signal between 3 and 5dpi (%PfApiAT2-smHA positive 3dpi vs 4dpi *p*<0.0001, 4dpi vs 5dpi *p*=0.0004, Kruskal-Wallis with post-hoc Dunn’s correction). Close to half of PfApiAT2-smHA oocysts per midgut were smHA-positive at 4dpi. Each data point represents % oocysts per midgut microscopically assessed as positive for PfApiAT2-smHA signal, and line represents median, from 3 pooled biological replicates. Only oocysts stained positive for anti-GAPDH and guts with more than 1 oocyst were considered for quantification. Data shown represents median +/- 95% confidence intervals of parasites for each gut assessed microscopically, from 3 pooled biological replicates. Different letters indicate statistically significant differences, details of statistical tests are provided in Supplementary Data. (d) PfApiAT2-smHA oocysts at day 7 post infection, co-stained for (i) HA and (ii) the capsule protein Cap380. (iv) smHA-signal can be observed under the capsule, suggesting PfApiAT2 localization to be the plasma membrane. Representative images from two independent biological replicates. Source data are provided as a Source Data file.

Indirect immunofluorescence assay (IFA) analysis in asexual blood-stage PfApiAT2-smHA parasites revealed membrane localization of PfApiAT2 consistent with previously published data (Supplementary Figure 4C) [28]. We infected *An. gambiae* with PfApiAT2-smHA parasites and performed IFAs in early oocysts, which revealed increasing peripheral localization over 3-5dpi, with percent oocysts observed to be PfApiAT2-smHA positive per midgut significantly increasing each day between 3dpi and 5dpi (percent PfApiAT2-smHA positive 3dpi vs 4dpi *p*<0.0001, 4dpi vs 5dpi *p*=0.0004). 56% of oocysts/midgut expressed PfApiAT2-smHA at 4dpi (Figure 3B, C). To precisely localize PfApiAT2 on the oocyst surface, we co-stained midguts with an antibody against the capsule marker Cap380 at 7dpi, revealing a clear separation of PfApiAT2-smHA from the capsule (Figure 3D). These data demonstrate that PfApiAT2 is localized on the plasma membrane and its expression coincides with the onset of the growth defects observed in PfApiAT2-KO parasites.

### PfApiAT2-KO oocysts stall and degenerate over time

In order to observe the morphological consequences of genetic ablation of *pfapiat2*, we performed transmission electron microscopy (TEM) of midguts infected with PfApiAT2-KO or WT parasites during early (7dpi, Figure 4A, B) and late (14dpi, Figure 4C-F) development. At 7dpi, no gross morphological defects other than a smaller size were observed in PfApiAT2-KO oocysts. We detected a typical distribution of mitochondria, nuclear mass and other membrane-bound organelles, and identified the crystalloid, associated with early oocyst growth in *P. berghei* (Figure 4B)[29, 30]. Later in development, at 14dpi, we instead observed that while WT oocysts were undergoing segmentation, with clearly distinguishable daughter sporozoites (Figure 4C, D), the majority of PfApiAT2-KO oocysts (77.8%) appeared degenerate, with an electro-lucent cytoplasm and no distinguishable membrane-bound organelles as well as a lack of nuclear material and a thickened capsule (Figure 4E, N=18 oocysts imaged). Interestingly, 22.2% of observed PfApiAT2-KO oocysts were not degenerate but appeared morphologically younger in development, lacking daughter sporozoites and containing multiple nuclei and membrane-bound organelles, suggesting they were stalled well before entering segmentation (Figure 4F). This suggests that while oocysts lacking PfApiAT2 appear to be able to stall for at least 7 days, most are unable to maintain this stalled state, and eventually degenerate.

**Figure 4:**
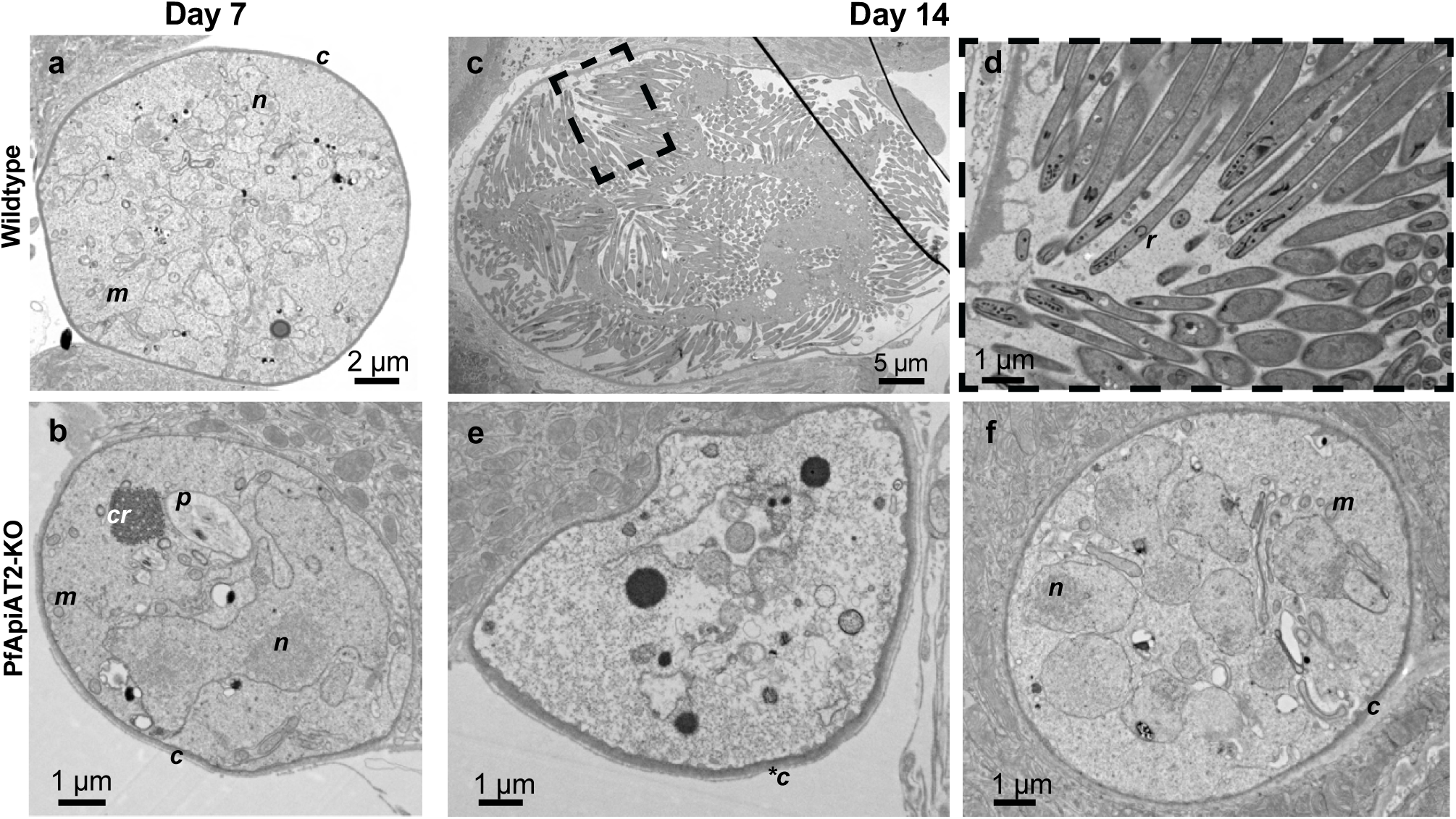
Ultrastructural analysis of PfApiAT2-KO reveals oocyst stalling and late-onset degeneracy. (a, b) At day 7 post infection, no gross morphological differences are evident between WT (a) and PfApiAT2-KO (b) oocysts. To note, organellar structures that can be observed include the capsule (*c*), the nucleus (*n*), the mitochondria (*m*). PfApiAT2-KO oocysts also show presence of the hemozoin crystals in a malaria pigment vacuole (*p*) left over from growth in the blood-stage, and the crystalloid (*cr*), a temporary organelle associated with young oocysts in *P. berghei* development [29]. Later in development, while WT oocysts (c, d) show signs of segmentation and the end of sporogony, with budding daughter sporozoites containing rhoptries (*r*), PfApiAT2-KO oocysts can be observed either degenerate (e) with an electro-lucent cytoplasm and a thickened capsule (*\*c*), or stalled (f), with membrane bound organelles akin to earlier oocysts. Representative images from two independent transmission experiments.

### PfApiAT2 is a proline-specific transporter

Past work had shown that asexual blood-stage parasites lacking PfApiAT2 accumulate proline, suggesting a specificity of this amino acid as a substrate for this transporter [21]. As the transport function of PfApiAT2 cannot be tested in oocysts due to the limited parasite biomass compared to mosquito cells in an infected midgut, we utilized *Xenopus laevis* oocytes as a heterologous expression system. Codon-harmonized sequences of PfApiAT2 and PfApiAT2 C-terminally linked to an mNeonGreen fusion tag (PfApiAT2-mNG) were cloned into expression vectors and used to generate complementary RNA (cRNA) (Supplementary Figure 5A, Supplementary Table 1) [31]. *X. laevis* oocytes were surgically isolated, dissociated, defolliculated and injected with PfApiAT2-mNG cRNA. Expression of PfApiAT2-mNG was visualized via confocal microscopy 4 days post-injection, colocalizing with the plasma membrane, stained with membrane dye FM4-64 (Supplementary Figure 4B), confirming correct membrane localization.

To test proline flux across the membrane, *Xenopus* oocytes injected with cRNA encoding PfApiAT2 and PfApiAT2-mNG were incubated in flux buffer containing ^14^C-proline. Oocytes expressing both injected constructs showed an accumulation of ^14^C-proline significantly greater than non-injected oocytes (Figure 5A, non-injected vs PfApiAT2-mNG injected *p*<0.0226, non-injected vs PfApiAT2 *p*<0.016, one-way ANOVA with post hoc Tukey’s multiple comparison). We performed flux assays across a range of ^14^C-proline concentrations and observed that PfApiAT2 and PfApiAT2-mNG mediated proline transport conformed to first-order Michaelis-Menten kinetics (K_m_ and V_max-apparent_ and traces of proline acquisition in Figure 5B).

**Figure 5:**
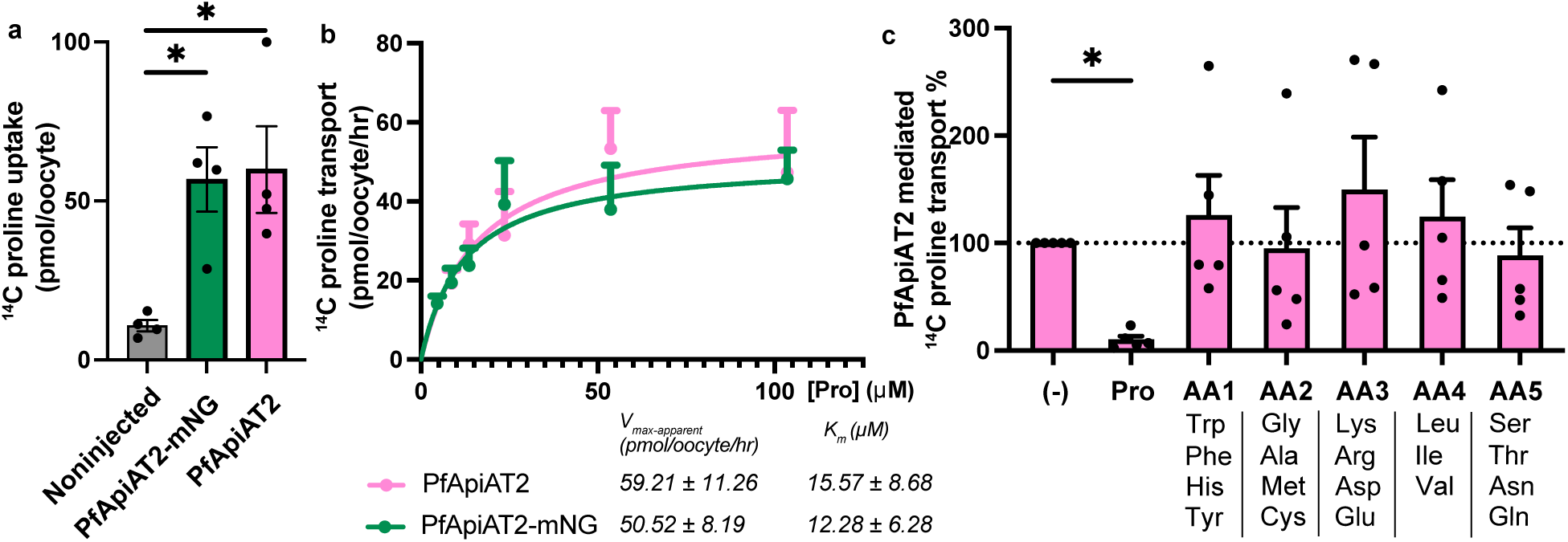
Heterologous expression of PfApiAT2 in *X. laevis* oocytes reveals its function as a proline-specific transporter. (a) *Xenopus laevis* oocytes injected with RNA coding for a fusion protein consisting of PfApiAT2 C-terminally tagged with fluorescent protein mNeonGreen (ApiAT2-mNG) or untagged PfApiAT2, when incubated in media containing 100 μM ^14^C-proline, show accumulation of radioactivity significantly greater than non-injected oocytes (Noninjected vs PfApiAT2-mNG *p*=0.016, noninjected vs PfApiAT2 *p*=0.016, One-way ANOVA, with post-hoc Tukey’s multiple comparison correction). Data shown is means +/- SEM of 1 hour of flux from 4 biological replicates of 7-10 oocytes per biological replicate. (b) Oocytes expressing both tagged and untagged PfApiAT2 take up ^14^C-proline from surrounding media with first-order kinetics of transport. Data shown is pmol of ^14^C-proline taken up by oocytes, normalized to uptake by non-injected controls, means +/- SEM from 4 biological replicates, containing 7-10 oocytes per biological replicate, and includes the data shown in (a). Kinetics parameters derived from kinetics curve. (c) Substrate competition experiments show PfApiAT2 is a proline specific transporter. Uptake of 5μM 1μCi/ml ^14^C-proline cannot be inhibited by any other non-radiolabeled substrate other than unlabeled proline. Competitors (other than proline) were pooled into groups of three or four, each competitor amino acid at 500μM (100-fold molar excess compared to radiolabeled proline). ((-) vs Pro *p*=0.0372; (-) vs AA1 *p*=0.8767; (-) vs AA2 *p*=0.9015; (-) vs AA3 *p*=0.4595; (-) vs AA4 *p*=0.8767; (-) vs AA5 *p*=0.9015, repeated measures one-way ANOVA, post-hoc Holm-Šídák’s multiple comparisons correction). Data reported is means +/- SEM, 5 biological replicates, 7-10 oocytes per biological replicate, details of statistical tests provided in Supplementary Data. Source data are provided as a Source Data file.

In order to understand substrate specificity, ^14^C-proline transport assays were performed in the presence of non-radiolabeled competitor amino acids, pooled into groups based on physical and chemical properties at 100x molar excess compared to ^14^C-proline (Figure 5C). Only non-radiolabeled proline was able to compete the uptake of ^14^C-proline, while all other competitor amino acid groups showed no significant effect ((-) vs Pro *p*=0.0372, (-) vs all other groups non-significant, repeated measures one-way ANOVA with post hoc Holm-Šídák’s multiple comparisons, Supplementary Data for all other tests).

To test whether the transport of proline by PfApiAT2 requires co-transported ions, we used a two-electrode voltage clamp (TEVC) setup to measure membrane potential during proline flux. We observed that proline uptake was found to be electroneutral when probed with a positive or negative potential across the membrane (Supplementary Figure 5C). Structural prediction of PfApiAT2 using AlphaFold revealed an electropositive bias across the membrane, suggestive of a membrane protein that has strongly conserved topology of insertion into canonically charged plasma membranes [32]. A channel with both electro-positive and electro-negative residues was predicted, consistent with electroneutral proline transport given this amino acid is polar and neutral (Supplementary Figure 5D).

### PfApiAT2-KO oocyst growth is rescued by supplementation of exogenous nutrients

Given PfApiAT2’s role in transporting proline, and the observed stalling of early oocyst growth, we hypothesized that PfApiAT2-KO oocysts experience a nutritional deficit. To test this hypothesis, we provided mosquitoes previously infected with either WT or PfApiAT2-KO parasites with an additional non-infectious blood meal to determine whether oocyst growth could be rescued by supplementary nutrients. We compared oocyst size and sporozoite rates in infected mosquitoes blood fed twice (2BF group) versus infected mosquitoes blood fed only at the time of infection (1BF group). A third group was instead provided a proteinaceous meal (fatty acid-free bovine serum albumin, BSA group) at the time of the second blood feeding, which has been previously described to increase oocyst growth [5, 7] (Figure 6A). In NF54 WT-infected controls, at 7dpi, an additional blood meal increased oocyst size while the BSA meal had a marginal but not statistically significant effect (1BF vs 2BF *p*=0.002, 1BF vs BSA *p*=0.7435, Kruskal-Wallis). Interestingly, both the blood and protein meal partially rescued the size of PfApiAT2-KO oocysts (Figure 6B) (1BF vs 2BF *p*=0.0013, 1BF vs BSA *p*=0.0007, Kruskal-Wallis). When we compared the size of all individual oocysts from the three groups (rather than mean size per mosquito, as done normally), we found that every PfApiAT2-KO oocyst appeared to have resumed growth upon nutrient supplementation (Supplementary Figure 6A). Consistent with these results, salivary gland sporozoite numbers were also partially rescued at 14dpi (Figure 6C) (percent infected mosquitoes PfApiAT2KO 1BF vs 2BF *p*<0.0001, 1BF vs BSA *p*=0.0035, Kruskal-Wallis).

**Figure 6:**
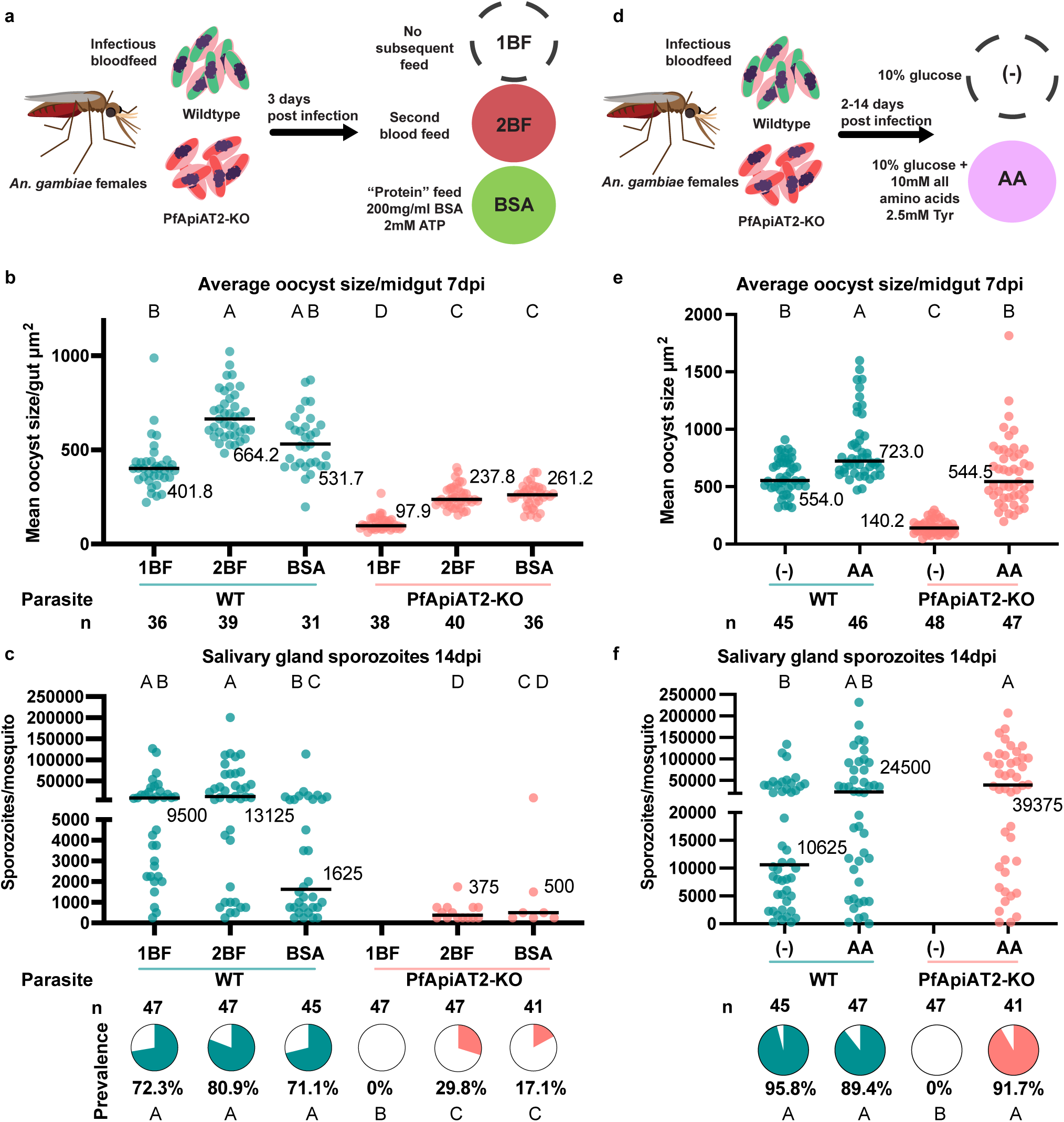
PfApiAT2-KO oocysts can be rescued with exogenous nutrients. (a) Scheme of multiple blood-feeding experiments. Female *An. gambiae* mosquitoes were infected with PfApiAT2-KO and WT parasites. Infected mosquitoes were either maintained as normal (1BF) or allowed to feed on a second non-infectious bloodmeal (2BF) or a protein meal consisting fatty-acid free bovine serum albumin (BSA), 3 days post-infection. (b) A second blood feed or protein feed partially rescued PfApiAT2-KO oocyst size (PfApiAT2-KO 1BF vs PfApiAT2-KO 2BF *p*=0.0013; PfApiAT2-KO 1BF vs PfApiAT2-KO BSA *p*=0.0007; WT 1BF vs PfApiAT2-KO 1BF *p*<0.0001; WT 2BF vs PfApiAT2-KO 2BF *p*<0.0001; WT BSA vs PfApiAT2 BSA *p*<0.0001). (c) Salivary gland sporozoite burden in mosquitoes infected with PfApiAT2-KO was also partially rescued with a second blood feed or protein feed (% infected mosquitoes PfApiAT2-KO 1BF vs PfApiAT2-KO 2BF *p*<0.0001; PfApiAT2-KO 1BF vs PfApiAT2-KO BSA *p*=0.0035; WT 1BF vs PfApiAT2-KO 1BF *p*<0.0001; WT 2BF vs PfApiAT2-KO 2BF *p*<0.0001; WT BSA vs PfApiAT2 BSA *p*<0.0001). (d) Scheme of sugar supplementation experiments. Infected mosquitoes were maintained on either normal diet comprising 10% glucose dissolved in water (-), or 10% glucose supplemented with a mix of all amino acids (AA). (e) *Ad libitum* supplementation of amino acids completely rescued PfApiAT2-KO oocyst size at 7dpi (WT (-) vs PfApiAT2-KO (-) *p*<0.0001; PfApiAT2-KO (-) vs PfApiAT2-KO AA *p*<0.0001; WT (-) vs PfApiAT2-KO AA *p*>0.9999). (f) Salivary gland sporozoite burden of mosquitoes fed PfApiAT2-KO parasites was completely rescued by supplementation of all amino acids (% infected mosquitoes WT (-) vs PfApiAT2-KO (-) *p*<0.0001; PfApiAT2-KO (-) vs PfApiAT2-KO AA *p*<0.0001; WT (-) vs PfApiAT2-KO AA *p*>0.6773). For size data (b, e), each dot represents average area of oocysts per midgut from midguts with > 2 midguts, and line represents median, data assessed using Kruskal-Wallis test with post-hoc Dunn’s correction. For sporozoite data, (c, f) each dot represents sporozoites counted via hemocytometer per dissected mosquito, and line represents median, assessed using Kruskal-Wallis test with post-hoc Dunn’s correction, and pie charts represent % mosquitoes infected, assessed using two-tailed Fisher’s Exact test. Data is pooled from 3 independent biological replicates. Different letters indicate statistically significant differences, details of statistical tests and all comparisons provided in Supplementary Data. Source data are provided as a Source Data file.

We hypothesized that direct supplementation of amino acids may also be able to rescue oocyst growth in PfApiAT2-KO. Therefore, soon after infection (2dpi) with WT or PfApiAT2-KO parasites, mosquitoes were maintained either on standard 10% glucose solution ((-) group) or on the same solution supplemented with a mixture of all amino acids (AA group) (each amino acid at 10mM except tyrosine, provided at 2.5mM due to solubility limits) for the duration of infection (Figure 6D). Amino acid supplementation increased the size of WT parasites ((-) vs AA *p*=0.0246, Kruskal-Wallis) and completely rescued the growth of PfApiAT2-KO oocysts at 7dpi ((-) vs AA *p*<0.0001, Kruskal-Wallis) (Figure 6E). Salivary gland sporozoite numbers and prevalence in PfApiAT2-KO at 14dpi were also fully rescued by amino acid supplementation (Figure 6F, percent infected mosquitoes PfApiAT2-KO (-) vs AA *p*<0.0001 two-tailed Fisher’s Exact). Notably, supplementing the sugar of infected mosquitoes with a 20mM proline solution from the time of infection did not increase the size of PfApiAT2-KO oocysts at 7dpi (Supplementary Figure 6B), suggesting PfApiAT2 may be the sole proline transporter in growing oocysts. Further investigation of the compartmentalization of sugar-meal derived amino acids into mosquito microenvironments accessible to the parasite is warranted.

Overall, these data show that PfApiAT2-KO oocysts are rescued by nutrients derived from either an additional blood meal, a protein-rich BSA-meal or amino acids. These data hint at oocysts being capable of synthesizing proline from other amino acids, a process that has been demonstrated in asexual blood-stage parasites, but further investigation is needed to confirm this hypothesis [21].

## Discussion

In this study, we determined that PfApiAT2 is an amino acid transporter essential for *P. falciparum* development through *An. gambiae* mosquitoes. Loss of PfApiAT2 in asexual blood-stages is implicated in resistance to halofuginone; we found that SNPs in the *pfapiat2* locus conferring HFG resistance induce a severe fitness cost during oocyst growth and result in a dramatic reduction of salivary gland sporozoites. These results were fully recapitulated by genetic ablation of the *pfapiat2* locus, revealing that PfApiAT2 is essential for oocyst growth. PfApiAT2-KO parasites formed stalled, stunted oocysts unable to develop sporozoites for onward transmission, and underwent degeneracy. Additionally, we determined that PfApiAT2 localizes to the plasma membrane of growing oocysts between day 3 and 5 (the time at which PfApiAT2-KO oocyst growth stalls), right under the parasite capsule, which suggests the capsule is permeable to amino acids like proline, in line with observations in *P. berghei* of a capsule permeable to water and small solutes [33]. Using heterologous expression in *X. laevis* oocytes, we demonstrated that PfApiAT2 is capable of proline-specific transport across the membrane. We further determined that PfApiAT2-KO oocyst growth can be partially or fully rescued by providing mosquitoes with supplemental nutrients in the form of an additional bloodmeal, BSA or amino acid mixtures.

Interestingly, in the asexual blood-stage, this transporter maintains proline homeostasis but is not necessary for parasite growth [21], likely due to the vast quantity of intracellular hemoglobin-derived proline, internalized from the host red blood cell. The essential function of PfApiAT2 during oocyst development may instead be due to the high abundance of free proline in the mosquito microenvironment following a blood meal, as shown in *Aedes aegypti* where proline is postulated to act as a sink for reactive nitrogen species and an energy source in powering flight, alongside its function as a proteogenic amino acid [34, 35]. Moreover, in *Anopheles stephensi* mosquitoes, where proline levels also increase after an infectious or non-infectious blood-feeding, hemolymph proline decreases at a faster rate in *P. berghei* infected mosquitoes than in uninfected ones, compatible with a demand for this amino acid by growing parasites [36].

Mosquitoes digest their blood meals within two to three days and utilize blood-derived nutrients for oogenesis, and excess nutrients that are not taken up by the developing eggs may be utilized by the parasite for growth [7, 37–39]. While single-cell RNA sequencing of *P. falciparum* during mosquito stages suggests that PfApiAT2 is expressed from 2dpi onwards [40], we only detected tagged protein at the plasma membrane around 4dpi, i.e. 2 days after oocysts have started forming. This timing of expression suggests that PfApiAT2 is necessary for the growth of parasites under baseline hemolymph metabolite conditions, once mosquitoes have digested their blood meal. Consistently, PfApiAT2-KO parasites showed no defect in establishing infection, and we were able to rescue oocyst growth after providing additional nutrients, including *ad libitum* amino acid supplementation. We also observed that PfApiAT2-KO oocysts are morphologically normal up until at least 7dpi but show aberrations by 14dpi, suggesting they can withstand starvation for a few days, likely by entering a stalled state from which they can be rescued by additional nutrients, as has been previously suggested for *P. berghei* [8]. In asexual blood-stage parasites, *Plasmodium* can synthesize proline from arginine via ornithine, and this pathway contributes to dysregulation of proline homeostasis in asexual stage PfApiAT2-KO parasites [2, 21, 41]. We hypothesize this is the pathway through which PfApiAT2-KO oocysts are rescued under nutrient supplementation conditions (but not by proline alone), a hypothesis supported by the detection of transcripts of enzymes in this pathway during oocyst development [21, 40]. Alternatively, oocysts may express other low-affinity proline transporters that facilitate PfApiAT2-KO rescue during supplementation. While we cannot rule out that PfApiAT2-KO oocysts may experience some other form of nutritional stress, our data strongly implicates an inability to scavenge proline from the mosquito host as driving the growth defect.

HFG-resistant parasites exhibited a strong fitness cost associated with mutations in the *pfapiat2* locus during oocyst development, suggesting this locus is non-permissive to non-synonymous mutations. Consistent with this finding, few non-synonymous mutations and a lower non-synonymous/synonymous mutation ratio than the median across the genome are found in *pfapiat2* in whole genome sequencing of clinical *P. falciparum* samples (Supplementary Figure 7). Despite the non-essentiality during blood-stage growth, parasites are likely to be under strong purifying selection at this locus due to its key function during transmission through the mosquito vector. While the blood-stage dispensability of PfApiAT2 explains rapid recrudescence of parasites treated with HFG precursors [42], which has resulted in reduced interest in pursuing proline-competitive cPRS compounds [19], our findings suggest that HFG-R parasites with mutations in the *pfapiat2* locus will be severely impaired in their ability to be transmitted by mosquitoes, thereby limiting their spread.

Recent work from our lab has shown that *Plasmodium* parasites can be efficiently blocked during their development in the *Anopheles* female using antimalarials provided via mosquito nets or other surfaces [43, 44]. Our data uncovering proline homeostasis as critical for oocyst growth suggest factors essential for nutrient scavenging by the parasite in the mosquito vector may be promising druggable candidates for transmission-blocking strategies, expanding the malaria control toolbox.

## Methods

### Routine culture of asexual stage *Plasmodium falciparum*

*P. falciparum* NF54 strain (BEI Resources, MRA-1000) was cultured as asexual stages at 37°C between 0.2 and 2% parasitemia in human erythrocytes at 5% hematocrit (Research Blood Components, Watertown, MA) using RPMI medium 1640 supplemented with 25mM HEPES, 10mg/l hypoxanthine, 0.2% sodium bicarbonate, and 10% heat-inactivated human serum (Innovative Research, Novi, MI, and BioIVT, Woodbury, NY) under a gas mixture of 5% O_2_, 5% CO_2_, balanced N_2_.

### Generation of HFG-resistant parasites

Asexual stage parasites were cultured in the presence of 3nM HFG for 6 days until parasites could no longer be observed by thin smear. Recrudescent parasites were observed between 15-20 days following removal of drug. DNA was extracted from asexual stage parasites using Qiagen Blood Mini Kit and the *pfapiat2* locus was amplified via PCR (primers P1 and P9, Supplementary Table 1). Bulk parasite culture was genotyped via Sanger sequencing (Azenta/Genwiz). R345I blood-stage culture was genotyped by ligating the amplified *pfapiat2* locus into a plasmid using Zero Blunt Topo PCR Cloning Kit (Thermo Fisher) and cloned and amplified in NEB DH5α *E. coli* due to issues with recovering reliable Sanger sequencing from PCR product. Extracted DNA was genotyped using longread sequencing (Plasmidsaurus). Age-matched parental wild-type parasites were used as control for drug susceptibility and transmission assays.

### Generation of transgenic PfApiAT2-KO parasites

Homology-directed repair using CRISPR-Cas9 was used to insert a human-dihydrofolate (hDHFR) cassette into the *pfapiat2* locus as described previously [21]. Briefly, one plasmid encoding Cas9 enzyme and homologous regions (HR) upstream and downstream (donor), and two plasmids with guide sequences were generated and purified as described. Parasite culture pellet at 5% hematocrit and ∼5% parasitemia was washed in Cytomix media (120 mM KCl, 0.15 mM CaCl_2_, 2 mM EGTA, 5 mM MgCl_2_, 10 mM K_2_HPO_4_/KH_2_PO_4_, pH 7.6, 25 mM HEPES, pH 7.6) and mixed with the two guide plasmids and linearized donor DNA dissolved in Cytomix. The suspension was electroporated using a Bio-Rad Gene Pulser at 0.34 kV and 950 µF and allowed to recover in culture media with fresh red blood cells. 10-12 hours post electroporation, culture was washed and resuspended in media containing 5nM WR9921 to select for transgenic parasites. Genomic integration was confirmed via PCR of recrudescent parasites, which were maintained on WR9921 until induction of gametocytes. Age-matched parental wild-type parasites were used as control for subsequent assays.

### Generation of transgenic PfApiAT2-smHA parasites

The high-avidity smHA tag was inserted C-terminally into the *pfapiat2* genomic locus using CRISPR-Cas9. Two guide sequences were selected (using benchling.com) at the C-terminus of the genomic locus: (5’ AATAGGATGTATATATTTGA 3’) and (5’ AAAAAAAAAGAAGAAGCAAA 3’). These sequences were ligated into a linearized pDC2 backbone plasmid (a kind gift from Dr. Marcus Lee) [26] downstream of a U6 promoter, proximal to the invariant region of the chimeric sgRNA scaffold to generate the guide plasmids. Donor plasmid was made by PCR amplification and introduction of shield mutations into regions 5’ and 3’ of the guide sites from the genomic locus using primers P10-P13. These were ligated via Gibson assembly into a pJPM10 plasmid backbone (a kind gift from Dr. Jeffery D. Dvorin) containing a Cam promoter upstream of an hDHFR cassette for negative selection using WR9921, BglII and NcoI restriction cut sites were introduced between the two HR, and an smHA construct was ligated downstream of the 5’ HR. Transfection of asexual blood-culture and selection under WR9921 for transgenic parasites was performed as described above, and successful integration of the smHA tag to the C-terminus of PfApiAT2 was confirmed via PCR around the integration site.

### Assessment of asexual blood-stage HFG sensitivity *in vitro*

HFG sensitivity of drug-selected and transgenic lines was assessed and compared to WT using a SYBR Green I fluorescence assay as previously described[45]. Briefly, asexual blood-stage culture was synchronized using washes in 5% D-sorbitol and seeded at 1% ring stage culture, 1% hematocrit in black-walled clear-bottom 384 well plates containing 12-point dilution series of HFG dissolved in DMSO, with technical triplicates. Dilution series of dihydroartemisinin and atovaquone were included as controls to ensure any shifts in EC_50_ observed were specific to HFG. Compounds were dispensed with an HP D300 Digital Dispenser (Hewlett Packard, Palo Alto, CA, USA). Growth was assessed 72 hours post exposure to HFG via SYBR Green staining of parasite DNA. Fluorescence intensity measurements were performed on a SpectraMax M5 (Molecular Devices, Sunnyvale, CA, USA) and analyzed in GraphPad Prism (GraphPad Software, La Jolla, CA, USA) after background subtraction and normalization to control wells.

### Transmission of parasites to female *An. gambiae* mosquitoes

Stage V female and male gametocytes were induced by raising parasitemia over 4% and incubating cultures for ∼14–20 days with daily media change. For transmission experiments following drug selection or transgenic manipulation, cultures were not cloned, as long-term culture of blood-stage parasites has been associated with loss of gametocytogenesis and transmissibility to mosquitoes [46].

Colony of *An. gambiae* G3 mosquitoes was reared as previously described [7]. Cages of adult females isolated 3-7 days post-eclosion maintained on 10% sugar solution, were blood-fed on a mixture of mature stage V *P. falciparum* gametocytes, red blood cells and serum (40% hematocrit, 1:3 ratio of gametocyte culture and packed red blood cells) via a standard membrane feeding assay for 60min in a custom-built glovebox (Process Control Solutions, Shrewsbury, MA). Females not completely engorged were removed via aspiration into 80% ethanol and fed mosquitoes were maintained on 10% sugar solution-soaked cotton pads. At dissection timepoints, infected mosquitoes were aspirated into 80% ethanol and transferred to cold 1X PBS.

### Assessment of oocyst development

Midguts from infected midguts were dissected in PBS and stained in 2mg/ml mercury dibromofluorescein disodium salt solution (mercurochrome, Sigma-Aldrich) in ddH_2_O for 14-17 minutes at room temperature. Guts were mounted on yellow 10-chamber slides in 4μl 0.2mg/ml mercurochrome solution and covered with a coverslip. Samples were imaged using an inverted light microscope (Olympus Inverted CKX41 microscope) at 100X magnification. Images were also taken at 200X and 400X magnification to capture smaller oocysts.

Images were stitched using a Fiji macro, and oocysts counted and sized using a machine learning algorithm developed in the lab, OocystMeter, and confirmed manually [47]. Oocysts too small to be measured by the software were sized manually using Fiji [48]. Oocyst counts per midgut from each midgut containing >1 oocysts, prevalence of infection (percent midguts where ≥1 oocysts present), and mean oocyst size per midgut from each mosquito are shown. For scatter plots, median is shown, and statistical significance was determined via Mann-Whitney tests where two groups were compared, Kruskal-Wallis tests with multiple comparison corrections for more than two groups, and pairwise Chi-squared tests for prevalence. GraphPad Prism was used to plot the data and perform statistical tests, and details of statistical tests are provided in Supplementary Data.

### Assessment of salivary gland sporozoite numbers

Salivary glands from infected mosquitoes were dissected into 50μl PBS + 1% Penn/Strep on ice. Samples were homogenized using a micro-pestle. 10μl of suspension was loaded onto Neubauer hemocytometers and sporozoite numbers counted using an inverted phase-contrast microscope at 200X or 400X. Number of sporozoites per mosquito containing >1 sporozoite microscopically observed and prevalence of infection (percent mosquitoes with ≥1 sporozoite observed) were plotted. For scatter plots, median is shown, and statistical significance was determined via Mann-Whitney tests where two groups were compared, Kruskal-Wallis tests with multiple comparison corrections for more than two groups, and pairwise Chi-squared tests for prevalence. GraphPad Prism was used to plot the data and perform statistical tests, and details of statistical tests are provided in Supplementary Data.

### Multiple blood-feeding of infected mosquitoes

Adult mosquitoes were allowed to mate by housing males and females at a 2:1 ratio for 3 days pre-feeding. Female mosquitoes were isolated into cages prior to feeding and infected as described above. Cages contained an oviposition site (a petri dish filled with wet cotton, covered with a filter paper) and subsequent feeding behavior was encouraged by starving mosquitoes for 24 hours before a second feed. Infected mosquitoes were fed 3 days post infection on uninfected blood, or a protein-meal: 200mg/ml fatty acid free bovine serum albumin (BSA, Sigma Aldrich A6003), 2mM ATP and 2.5% v/v food-dye in 1X PBS[5]. Unfed mosquitoes were removed following the second feed and fed mosquitoes were maintained on 10% glucose until dissection.

### Amino acid supplementation of mosquito sugar meal

Adult female *An. gambiae* mosquitoes were infected as described earlier in the text. Infected mosquitoes were provided (via soaked cotton pads) 10% glucose supplemented with amino acid mix containing all amino acids at 10mM each (except tyrosine, at 2.5mM, the solubility limit), provided *ad libitum* from 2dpi onwards. For single supplementation with proline, 20mM of L-proline dissolved in 10% glucose was provided *ad libitum* from 1dpi onwards.

### Generation of anti-Cap380 antibodies

Polyclonal sera were raised by GenScript (GenScript, NJ) in two rabbits using the epitope previously described [49]. Briefly, a recombinant peptide consisting of residues 1954–2068 from the sequence of *Pf*Cap380 (PF3D7_0320400) and a 6XHis N-terminal sequence was expressed in a BL21 Star (DE3) bacterial expression system, and affinity purified. 2 rabbits were injected with the recombinant construct 3 times and antibodies purified from rabbit sera. Sera from both rabbits were pooled and used for IFAs.

### Immunofluorescence assay localization of PfApiAT2 in oocysts

Infected mosquito guts were dissected in PBS and fixed in 4% PFA in PBS at room temperature for 1 hour, followed by 3 washes in PBS for 5 minutes each. Samples were blocked and permeabilized in 3% BSA and 0.1% TritonX (blocking buffer) at 4°C. Primary antibodies (Supplementary Table 2) were dissolved in blocking buffer and samples incubated overnight at 4°C or 1 hour at room temp. After washing 3 times in blocking buffer, samples were incubated for 30 min at room temp with secondary antibodies (Supplementary Table 2) dissolved in blocking buffer. After washing 3 times with PBS and staining with Hoechst (for DNA), guts were mounted in VectaShield Mounting Media (Fisher Scientific, NC9265087) under #1.5 coverslips. For time-course of PfApiAT2-smHA membrane localization, guts were imaged on a Zeiss Inverted Observer Z1 at 630X and membrane localization positivity was assessed by eye. For each biological replicate, the same exposure and acquisition settings were used between timepoints. WT samples as well as PfApiAT2-smHA samples stained without primary antibody were imaged to confirm staining specificity. Percent of oocysts positive or negative for peripheral smHA signal for each timepoint per gut were represented as a stacked bar and statistical significance determined by Kruskal-Wallis test between percent smHA positive oocysts per gut across timepoints correcting for multiple comparisons. Data plotted and statistical tests performed using GraphPad Prism. Representative images were acquired using a confocal Zeiss LSM980 with an Airyscan detector.

Images visualizing PfApiAT2-smHA localization with Cap380 were acquired using the confocal Zeiss LSM980 with an Airyscan detector. Images were analyzed using Zen Blue 3.0 and Fiji [48].

### Immunofluorescence assay localization of PfApiAT2 in asexual blood-stages

Asexual blood-stage IFAs were performed as previously described [50]. Briefly, air dried thin smears of asexual blood-stage parasites were fixed in 4% PFA, and washed with 1XPBS three times. Smears were permeabilized in 0.1% Triton-X in PBS for 10 min and washed in 1XPBS three times. Smears were blocked in 3% BSA in PBS overnight at 4°C and stained with primary antibodies (Supplementary Table 2) dissolved in blocking buffer for 1 hour at room temp. Slides were washed 3 times in blocking buffer and stained with secondary antibodies (Supplementary Table 2) and wheat germ agglutinin (WGA 1 in 1000 from 2mg/ml stock, 29022-1, Biotium, to stain the RBC membrane) dissolved in blocking buffer for 30 min at room temp. Slides were washed 3 times with PBS, mounted in VectaShield Mounting Media under #1.5 coverslips as above, and imaged on a confocal Zeiss LSM980 with an Airyscan detector.

### Transmission electron microscopy

Samples were fixed overnight in a mixture of 1.25% formaldehyde, 2.5% glutaraldehyde, and 0.03% picric acid in 0.1 M sodium cacodylate buffer, pH 7.4. The fixed tissues were washed with 0.1M sodium cacodylate buffer and post-fixed with 1% osmium tetroxide/1.5% potassium ferrocyanide in water for 2 hours. Samples were then washed in a maleate buffer and post fixed in 1% uranyl acetate in maleate buffer for 1 hour. Tissues were then rinsed in water and dehydrated through a series of ethanol washes (50%, 70%, 95%, (2x) 100%) for 15 minutes per solution. Dehydrated tissues were put in propylene oxide for 5 minutes before they were infiltrated in epon mixed 1:1 with propylene oxide overnight at 4°C. Samples were polymerized in a 60°C oven in epon resin for 48 hours. They were then sectioned into 80nm thin sections and imaged on a JEOL 1200EX Transmission Electron Microscope equipped with an AMT 2k CCD camera.

### Heterologous expression of PfApiAT2 in *Xenopus laevis* oocytes

Codon harmonized sequences for PfApiAT2 and PfApiAT2-mNeonGreen (encoding fusion protein PfApiAT2 C-terminally tagged with mNeonGreen) (Supplementary Table 1), were designed [51] from *pfapiat2* sequence extracted from PlasmoDB (accession Pf3D7_0914700) [52]. Sequences were cloned into a custom pMAX+ vector with a bacterial origin of replication, Amp-resistance cassette and T7 promoter sequence (Genwiz). Codon harmonized sequences were inserted between a 5’ and 3’ *Xenopus* globin UTR. Constructs were purified and coding RNA (cRNA) was generated using an mMESSAGE mMACHINE™ T7 Transcription Kit (Thermo Fisher Scientific, #AM1344).

Oocytes were surgically extracted from *X. laevis* frogs then isolated and defolliculated by collagenase treatment. Isolated oocytes were transferred to an Oocyte Ringer’s solution supplemented with Ca^2^+ and penicillin/streptomycin antibiotics (OR2++). Oocytes were each selected and injected with 50nl of cRNA resuspended in RNAse-free water. Oocytes were incubated at 15-17°C in OR2++ in 6 well plates with media refreshed daily until use [53].

Oocytes injected with 5ng of cRNA, 4 days post injection were placed in chambered coverslips (ibidi #80826) and imaged using with Zeiss Airyscan LSM980 confocal microscope at 100X magnification to visualize surface expression of PfApiAT2-mNeonGreen tagged fusion protein. Oocytes for two-electrode voltage clamp (TEVC) experiments were prepared as described previously [31]. Briefly, electrodes were pulled from borosilicate glass capillaries using a Flaming-Brown micropipette puller (Sutter Instruments) and filled with a 3M KCl solution containing 1.5% agarose. Pipette resistances were between 0.2 and 1.0 MΩ. Currents were recorded using an OC-725D (Warner) amplifier. Oocytes were perfused with a gravity-driven apparatus with a physiological buffer containing 2mM KCl, 96mM NaCl, 1mM MgCl_2_, and 5mM HEPES, buffered to pH 7.4 with NaOH, and 5mM L-proline where indicated. Currents were evoked by a voltage ramp protocol, in which the voltage is set to -80mV and then +80mV for 50 ms increments, separated by a 150 ms voltage ramp from a holding voltage of 0 mV. Pulses were repeated every 1s.

### Transport assays and kinetics of ^14^C -Proline uptake

Oocytes were injected with 25ng of RNA and maintained in OR2++ at 15-17°C for 4 days before performing transport assays. Non-injected oocytes from the same frog were used as a control. ^14^C labelled proline (Moravek, #MC263) at 1μCi/ml in ND96 flux buffer ( 96 mM NaCl, 2 mM KCl, 1 mM MgCl_2_, 1.8 mM CaCl_2_, 5mM HEPES[53]) was mixed with unlabeled proline at desired dilutions. Healthy injected or non-injected oocytes were transferred to polystyrene tubes and washed twice in 3ml room temperature ND96, removing residual supernatant. Oocytes were suspended in 100μl of flux buffer at appropriate proline concentration and incubated for 1 hour at 27°C. Flux buffer was removed, and oocytes were washed in 3ml cold ND96 twice, before being placed in 96-well white-bottom transillumination plates, one oocyte per well. 10 unused oocytes were added to the final row, along with 5μl of flux buffer as a radioactivity standard. Oocytes were lysed in 30μl 10% SDS overnight and resuspended in 150μl MicroScint PS (Revvity, 6013631). Plates were read in a Perkin Elmer MicroBeta^2^ liquid scintillation counter. Mean and SEM of ^14^C-proline uptake from 4 biological replicates measured at 100μM proline transport are represented and one-way ANOVA performed for statistical significance (after confirming normality with Shapiro-Wilk test). Mean and SEM values of PfApiAT2 and PfApiAT2-mNG mediated ^14^C-proline transport was calculated by subtracting uptake by non-injected oocytes from those expressing PfApiAT2 and PfApiAT2-mNG. Data from 4 biological replicates of transport traces are represented and used to fit a non-linear regression of the Michaelis-Menten equation and used to calculate K_m_ and V_max-apparent_. Data plotted and analyzed using GraphPad Prism.

### Competition of proline transport with non-radioactive amino acid substrates

Oocytes were injected with 12.5ng of RNA and cultured in OR2++ at 15-17°C for 4 days before performing competition assays, using age-matched non-injected oocytes as control. ND96 with 1μCi/ml ^14^C-proline and 5μM total proline was mixed with unlabeled competitor amino acids in groups of three to four amino acids. Each competitor amino acid was diluted to 500μM. Groups were based on physical and chemical properties. Unlabeled proline and water were used as controls for transport inhibition. Transport assay was performed over 1 hour at 27°C as described above. Specific PfApiAT2-mediated ^14^C-proline transport was calculated by subtracting signal from non-injected oocytes and mean and SEM of signal normalized to positive control (transport without competition) per biological replicate are represented. Shapiro-Wilk test performed to confirm normality and one way ANOVA performed to compare each competitor group with positive control. Data were plotted and analyzed using GraphPad Prism.

### PfApiAT2 structural prediction and analysis

Coding sequence of PfApiAT2 was found from PlasmoDB [52] (PF3D7_0914700) and structural prediction was done via AlphaFold[54]. Protein structure visualization was done via Pymol using the APBS electrostatics plugin (The PyMOL Molecular Graphics System, Version 2.5.2, Schrödinger, LLC).

### Population diversity analysis of *pfapiat2* locus

Genomic SNPs and dN/dS (nonsynonymous to synonymous SNP ratio) across the *P. falciparum* genome were extracted from PlasmoDB[52] and plotted using R. Nonsynonymous SNPs in the *pfapiat2* locus were plotted using publicly available Pf8 data analysed on HaploAtlas[55].

## Data Availability

The data generated in this study are provided in the Source Data file.

## Acknowledgements

We thank Dr. L. E. de Vries and Dr. W. R. Shaw for insightful discussions, the Catteruccia laboratory insectary core (C. Wisniewski, O. Hulai, K. Bumpus, A. Stanton, K. Thornburg) for technical support and mosquito colony maintenance and the Harvard Medical School Electron Microscopy Facility for sample processing and access to the TEM.

## Author Contributions

M.K., D.F.W., F.C., S.B. conceived of the study. M.K. performed asexual blood-stage genetic manipulation and HFG-resistance selection. C.T. designed plasmids for PfApiAT2-smHA tagging. N.S. performed gametocyte culturing and infectious feeds for mosquito infection experiments. M.K. and J.K. performed infection experiments. M.K. performed IFAs and TEM analysis. M.K. and R.L.S. designed expression constructs for *X. laevis* expression system and radio-uptake assay experiments. R.C.K. performed *X. laevis* injections and TEVC experiments. M.K. performed radio-uptake assays and *X. laevis* oocyte microscopy. L.D.P. oversaw and facilitated *X. laevis* experimentation. M.K., D.F.W. and F.C. wrote the initial manuscript draft. M.K., R.C.K., R.L.S., D.F.W., F.C. and S.B. reviewed and edited the manuscript. All authors approved the final manuscript.

## References

1. WHO, World Malaria Report 2024 2024, Geneva: World Health Organization.

2. Krishnan, A. and D. Soldati-Favre, Amino Acid Metabolism in Apicomplexan Parasites. Metabolites, 2021. 11(2).

3. Atella, G.C., et al., The major insect lipoprotein is a lipid source to mosquito stages of malaria parasite. Acta Trop, 2009. 109(2): p. 159–62.

4. Graumans, W., et al., When Is a Plasmodium-Infected Mosquito an Infectious Mosquito? Trends Parasitol, 2020. 36(8): p. 705–716.

5. Kwon, H., et al., Additional Feeding Reveals Differences in Immune Recognition and Growth of Plasmodium Parasites in the Mosquito Host. mSphere, 2021. 6(2).

6. Nyasembe, V.O., et al., Adipokinetic hormone signaling in the malaria vector Anopheles gambiae facilitates Plasmodium falciparum sporogony. Commun Biol, 2023. 6(1): p. 171.

7. Shaw, W.R., et al., Multiple blood feeding in mosquitoes shortens the Plasmodium falciparum incubation period and increases malaria transmission potential. PLoS Pathog, 2020. 16(12): p. e1009131.

8. Habtewold, T., et al., Plasmodium oocysts respond with dormancy to crowding and nutritional stress. Sci Rep, 2021. 11(1): p. 3090.

9. Saeed, S., A.Z. Tremp, and J.T. Dessens, Plasmodium berghei oocysts possess fatty acid synthesis and scavenging routes. Sci Rep, 2023. 13(1): p. 12700.

10. van Schaijk, B.C., et al., Type II fatty acid biosynthesis is essential for Plasmodium falciparum sporozoite development in the midgut of Anopheles mosquitoes. Eukaryot Cell, 2014. 13(5): p. 550–9.

11. Costa, G., et al., Non-competitive resource exploitation within-mosquito shapes evolution of malaria virulence. bioRxiv, 2017: p. 149443.

12. Hofer, L.M., et al., Additional blood meals increase sporozoite infection in Anopheles mosquitoes but not Plasmodium falciparum genetic diversity. Sci Rep, 2024. 14(1): p. 17467.

13. Lampe, L., et al., Metabolic balancing by miR-276 shapes the mosquito reproductive cycle and Plasmodium falciparum development. Nat Commun, 2019. 10(1): p. 5634.

14. Martin, R.E., et al., The ’permeome’ of the malaria parasite: an overview of the membrane transport proteins of Plasmodium falciparum. Genome Biol, 2005. 6(3): p. R26.

15. Amambua-Ngwa, A., et al., Chloroquine resistance evolution in Plasmodium falciparum is mediated by the putative amino acid transporter AAT1. Nat Microbiol, 2023. 8(7): p. 1213–1226.

16. Shafik, S.H., et al., The natural function of the malaria parasite’s chloroquine resistance transporter. Nat Commun, 2020. 11(1): p. 3922.

17. Kenthirapalan, S., et al., Functional profiles of orphan membrane transporters in the life cycle of the malaria parasite. Nat Commun, 2016. 7: p. 10519.

18. Fagbami, L., et al., The Adaptive Proline Response in P. falciparum Is Independent of PfeIK1 and eIF2alpha Signaling. ACS Infect Dis, 2019. 5(4): p. 515–520.

19. Tye, M.A., et al., Elucidating the path to Plasmodium prolyl-tRNA synthetase inhibitors that overcome halofuginone resistance. Nat Commun, 2022. 13(1): p. 4976.

20. Herman, J.D., et al., A genomic and evolutionary approach reveals non-genetic drug resistance in malaria. Genome Biol, 2014. 15(11): p. 511.

21. Bopp, S., et al., Disruption of P. falciparum amino acid transporter elevates intracellular proline and induces resistance to Prolyl-tRNA synthetase inhibitors. Cell Chem Biol, 2025. 32(10): p. 1293–1302 e5.

22. Fairweather, S.J., et al., Coordinated action of multiple transporters in the acquisition of essential cationic amino acids by the intracellular parasite Toxoplasma gondii. PLoS Pathog, 2021. 17(8): p. e1009835.

23. Parker, K.E.R., et al., The tyrosine transporter of Toxoplasma gondii is a member of the newly defined apicomplexan amino acid transporter (ApiAT) family. PLoS Pathog, 2019. 15(2): p. e1007577.

24. Lee, W.J., et al., Characteristics of Plasmodium vivax apicomplexan amino acid transporter 8 (PvApiAT8) in the cationic amino acid transport. Sci Rep, 2025. 15(1): p. 4234.

25. Boisson, B., et al., The novel putative transporter NPT1 plays a critical role in early stages of Plasmodium berghei sexual development. Mol Microbiol, 2011. 81(5): p. 1343–57.

26. Ullah, I., et al., Artemisinin resistance mutations in Pfcoronin impede hemoglobin uptake. bioRxiv, 2024.

27. Viswanathan, S., et al., High-performance probes for light and electron microscopy. Nat Methods, 2015. 12(6): p. 568–76.

28. Wichers, J.S., et al., Characterization of Apicomplexan Amino Acid Transporters (ApiATs) in the Malaria Parasite Plasmodium falciparum. mSphere, 2021. 6(6): p. e0074321.

29. Dessens, J.T., et al., Malaria crystalloids: specialized structures for parasite transmission? Trends Parasitol, 2011. 27(3): p. 106–10.

30. Dessens, J.T., A.Z. Tremp, and S. Saeed, Crystalloids: Fascinating Parasite Organelles Essential for Malaria Transmission. Trends Parasitol, 2021. 37(7): p. 581–584.

31. Gada, K.D., et al., Optogenetic dephosphorylation of phosphatidylinositol 4,5 bisphosphate in Xenopus laevis oocytes. STAR Protoc, 2023. 4(1): p. 102003.

32. von Heijne, G., Control of topology and mode of assembly of a polytopic membrane protein by positively charged residues. Nature, 1989. 341(6241): p. 456–8.

33. Saeed, S., A.Z. Tremp, and J.T. Dessens, Plasmodium sporozoite excystation involves local breakdown of the oocyst capsule. Sci Rep, 2023. 13(1): p. 22222.

34. Goldstrohm, D.A., J.E. Pennington, and M.A. Wells, The role of hemolymph proline as a nitrogen sink during blood meal digestion by the mosquito Aedes aegypti. J Insect Physiol, 2003. 49(2): p. 115–21.

35. Scaraffia, P.Y. and M.A. Wells, Proline can be utilized as an energy substrate during flight of Aedes aegypti females. J Insect Physiol, 2003. 49(6): p. 591–601.

36. Mack, S.R., S. Samuels, and J.P. Vanderberg, Hemolymph of Anopheles stephensi from uninfected and Plasmodium berghei-infected mosquitoes. 2. Free amino acids. J Parasitol, 1979. 65(1): p. 130–6.

37. Baton, L.A. and L.C. Ranford-Cartwright, Ookinete destruction within the mosquito midgut lumen explains Anopheles albimanus refractoriness to Plasmodium falciparum (3D7A) oocyst infection. Int J Parasitol, 2012. 42(3): p. 249–58.

38. Billingsley, P.F., Blood digestion in the mosquito, Anopheles stephensi Liston (Diptera: Culicidae): partial characterization and post-feeding activity of midgut aminopeptidases. Arch Insect Biochem Physiol, 1990. 15(3): p. 149–63.

39. Werling, K., et al., Steroid Hormone Function Controls Non-competitive Plasmodium Development in Anopheles. Cell, 2019. 177(2): p. 315–325 e14.

40. Yan, Y., et al., Mapping Plasmodium transitions and interactions in the Anopheles female. bioRxiv, 2024.

41. Gafan, C., et al., Characterization of the ornithine aminotransferase from Plasmodium falciparum. Mol Biochem Parasitol, 2001. 118(1): p. 1–10.

42. Jiang, S., et al., Antimalarial activities and therapeutic properties of febrifugine analogs. Antimicrob Agents Chemother, 2005. 49(3): p. 1169–76.

43. Probst, A.S., et al., In vivo screen of Plasmodium targets for mosquito-based malaria control. Nature, 2025. 643(8072): p. 785–793.

44. Paton, D.G., et al., Exposing Anopheles mosquitoes to antimalarials blocks Plasmodium parasite transmission. Nature, 2019. 567(7747): p. 239–243.

45. Johnson, J.D., et al., Assessment and continued validation of the malaria SYBR green I-based fluorescence assay for use in malaria drug screening. Antimicrob Agents Chemother, 2007. 51(6): p. 1926–33.

46. Claessens, A., et al., Culture adaptation of malaria parasites selects for convergent loss-of-function mutants. Sci Rep, 2017. 7: p. 41303.

47. Peng, D., et al., OocystMeter, a machine-learning algorithm to count and measure Plasmodium oocysts, reveals clustering patterns in the Anopheles midgut. bioRxiv, 2025.

48. Schindelin, J., et al., Fiji: an open-source platform for biological-image analysis. Nat Methods, 2012. 9(7): p. 676–82.

49. Itsara, L.S., et al., PfCap380 as a marker for Plasmodium falciparum oocyst development in vivo and in vitro. Malar J, 2018. 17(1): p. 135.

50. Cepeda Diaz, A.K., et al., Essential function of alveolin PfIMC1g in the Plasmodium falciparum asexual blood stage. mBio, 2023. 14(5): p. e0150723.

51. Lechner, H. Codon Harmonizer. Available from: http://biocatalysis.uni-graz.at/sites/codonharmonizer.html.

52. Aurrecoechea, C., et al., PlasmoDB: a functional genomic database for malaria parasites. Nucleic Acids Res, 2009. 37(Database issue): p. D539–43.

53. Broer, S., Xenopus laevis Oocytes. Methods Mol Biol, 2010. 637: p. 295–310.

54. Jumper, J., et al., Highly accurate protein structure prediction with AlphaFold. Nature, 2021. 596(7873): p. 583–589.

55. Lee, C., et al., Pf-HaploAtlas: an interactive web app for spatiotemporal analysis of Plasmodium falciparum genes. Bioinformatics, 2024. 40(11).

