## Supplementary figures and images for "PfApiAT2 is a proline transporter essential for the transmission of *Plasmodium falciparum* by the mosquito vector"

### Supplementary Figure 1

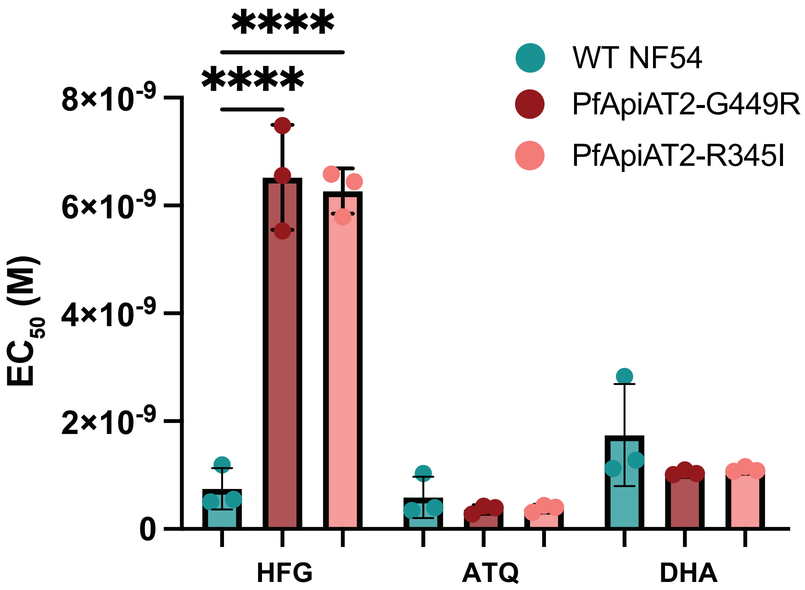

### Supplementary Figure 2

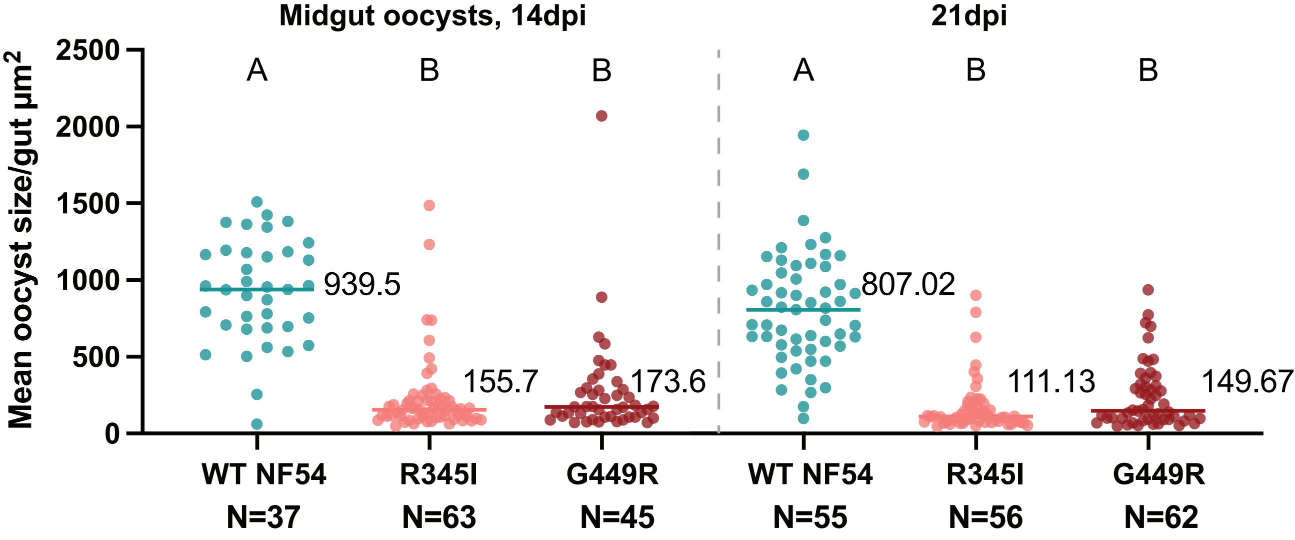

### Supplementary Figure 3

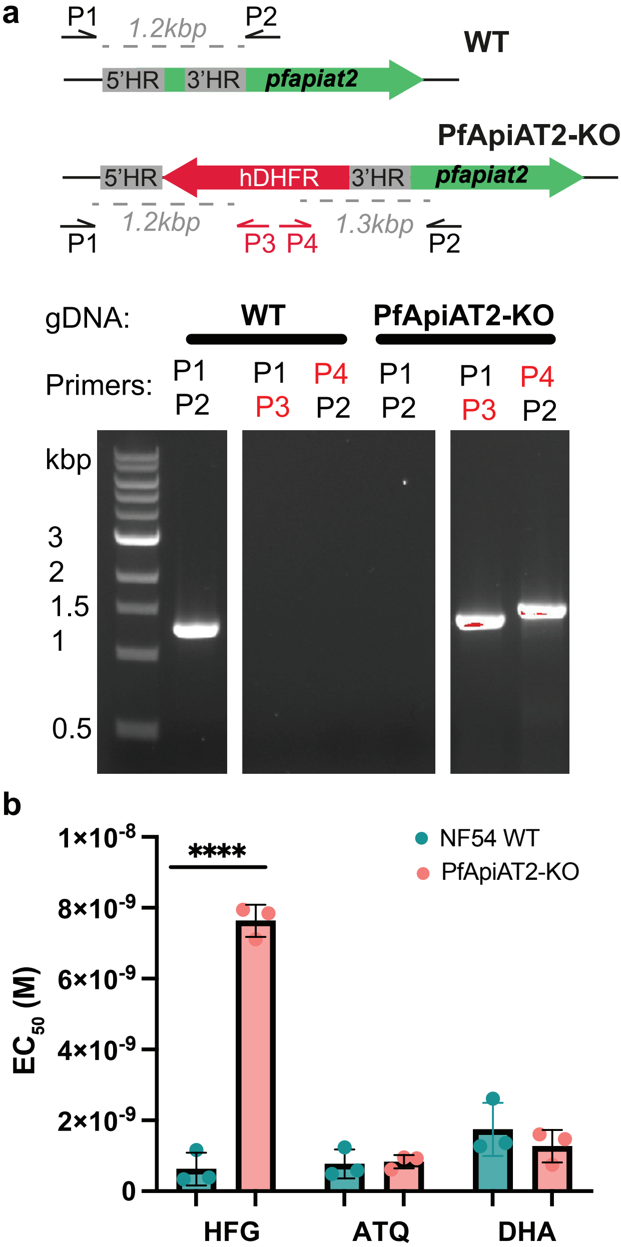

### Supplementary Figure 4

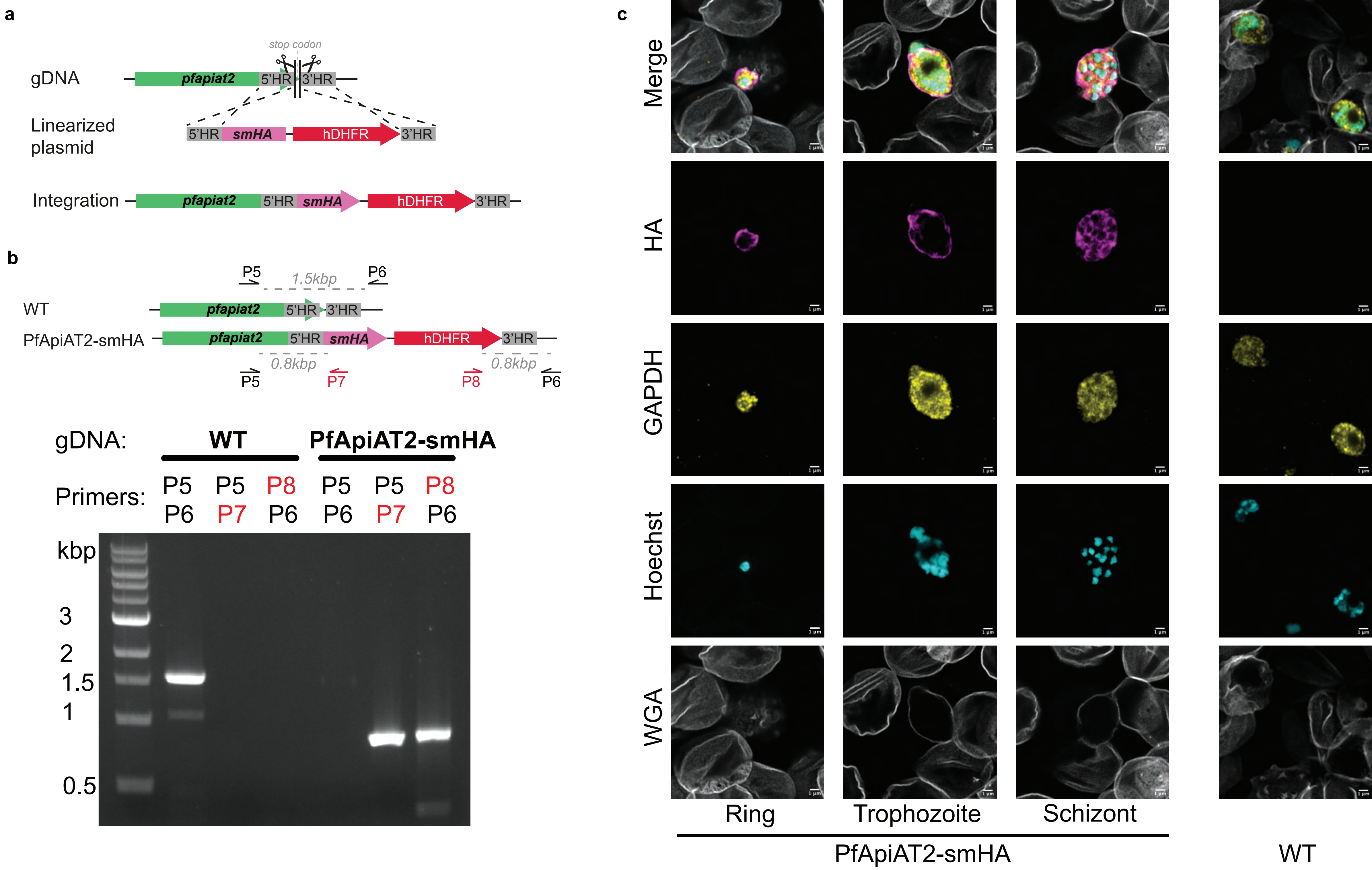

### Supplementary Figure 5

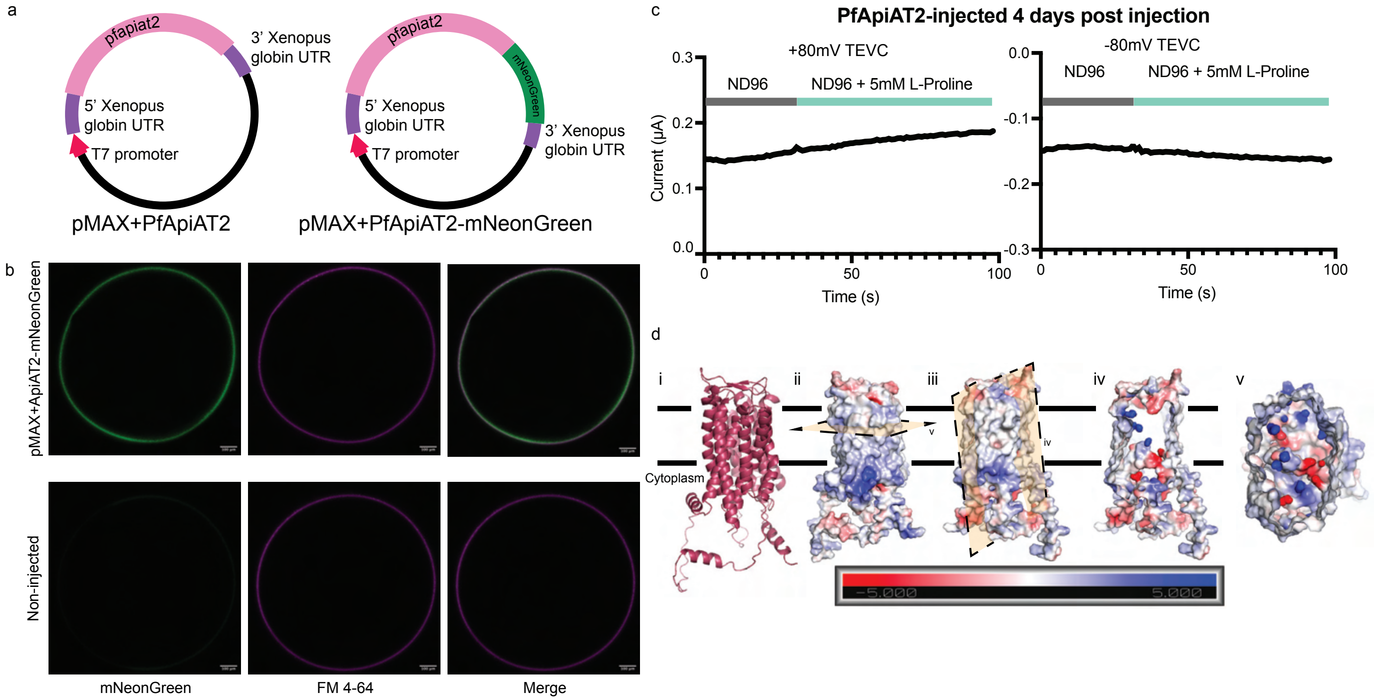

### Supplementary Figure 6

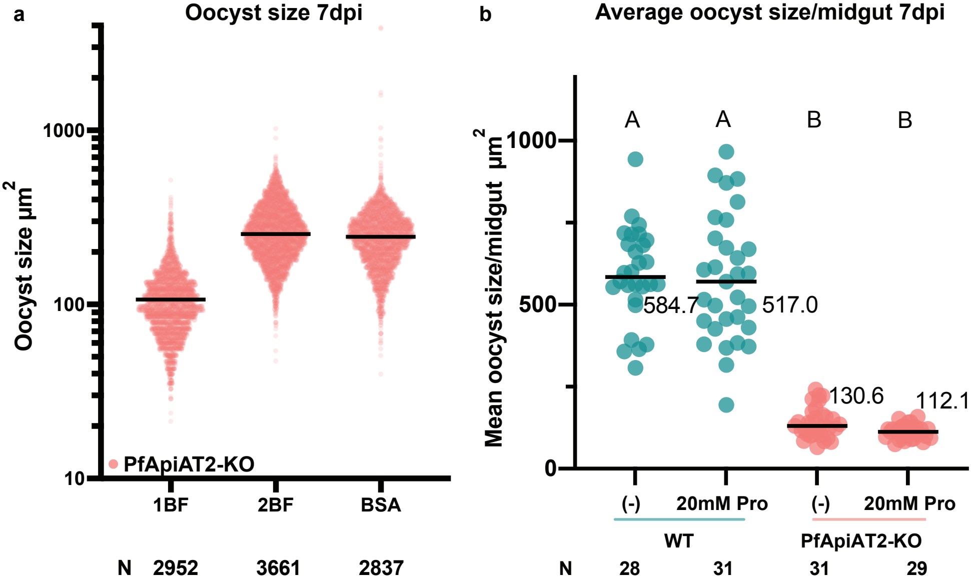

### Supplementary Figure 7

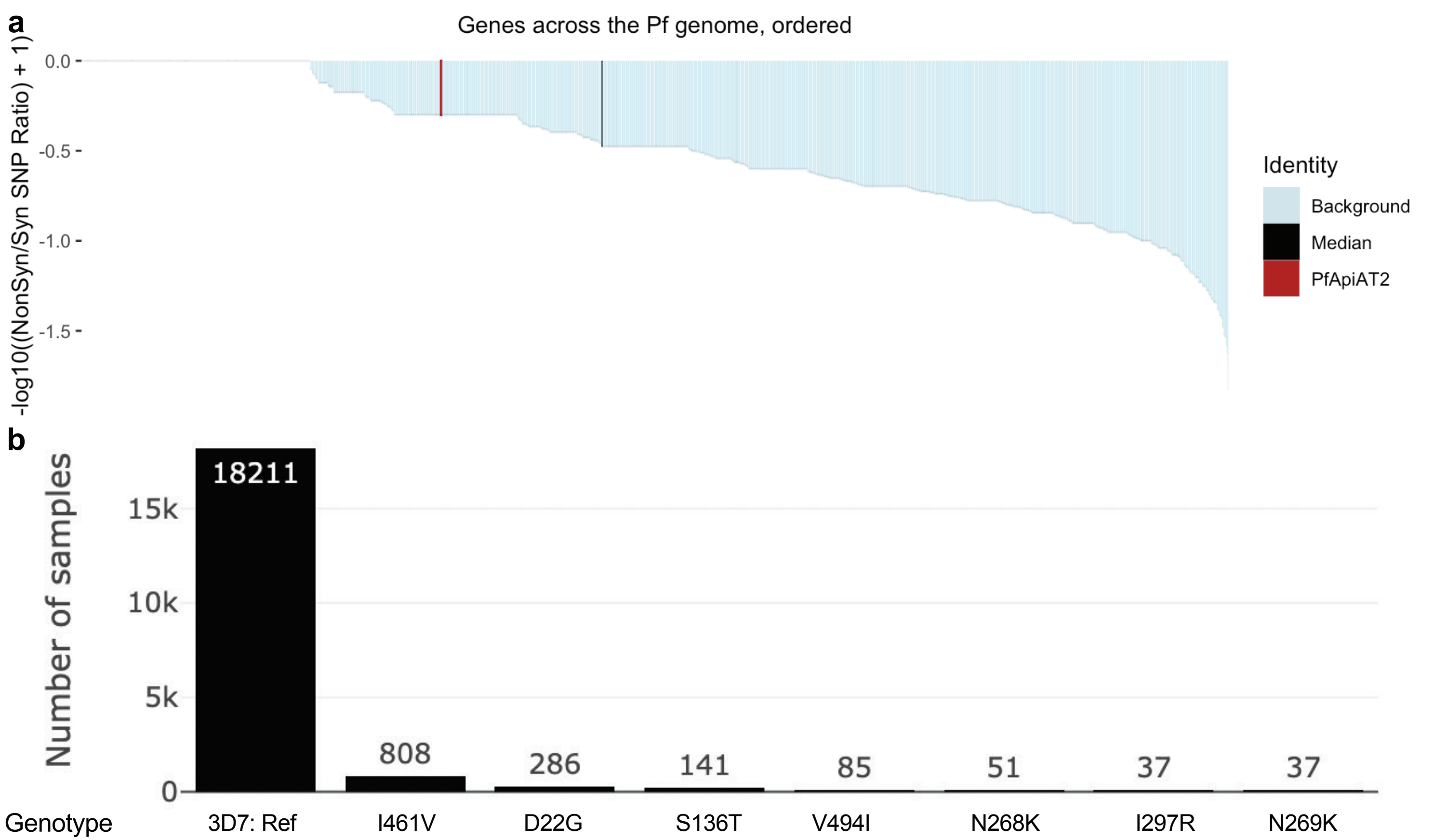
