## Supplementary Table 1 for "PfApiAT2 is a proline transporter essential for the transmission of *Plasmodium falciparum* by the mosquito vector"

| Sequence name | Sequence (5' --> 3') | Description |
|---|---|---|
| P1 | AAGGAACATATAGGATACCCAAC | PCR for PfApiAT2-KO insertion and sequencing HFG-R parasites |
| P2 | GAGGTTTAGATAATGAATTTCTGTCTG | PCR for PfApiAT2-KO insertion |
| P3 | GGTGTTCTCTCTGATGTCC | PCR for PfApiAT2-KO insertion |
| P4 | CTTTTATCATGCACATTGGAATAATAC | PCR for PfApiAT2-KO insertion |
| P5 | CAAATTAATAGTGAACGTGCTACC | PCR for PfApiAT2-smHA insertion |
| P6 | CCTTCCATATTAGGAACATTTATCATAAG | PCR for PfApiAT2-smHA insertion |
| P7 | CATCATAAGGGTAGCCAGCATAATC | PCR for PfApiAT2-smHA insertion |
| P8 | CAGGAAACAGCTATGACCATG | PCR for PfApiAT2-smHA insertion |
| P9 | GTTTGTATATATATGTGTGGGGGG | Sequencing HFG-R parassites |
| P10 | AGATCTCAATTGCCATGGGATCTTTCTTTATGACCAATGCATATG | Cloning 5'HR of PfApiAT2-smHA donor plasmid |
| P11 | GGATGTATATATTTGAAaGTTACAGAAAAGAAGAAAAACAAAAACCTCGAGATGTACCC | Cloning 5'HR of PfApiAT2-smHA donor plasmid |
| P12 | CGTCAGGATCATCGCGGCCGCGAAGCAAATGGTAAAACAATGCAG | Cloning 3'HR of PfApiAT2-smHA donor plasmid |
| P13 | CAGATGAATTTACTTCATTTTTCAAATATATAATGTGAGATCTCAATTGCCATGG | Cloning 3'HR of PfApiAT2-smHA donor plasmid |
| PfApiAT2_Xl | ATGGCGAGCGATGTTAGCAAGGAAAATATTAGCCAGCTAAAGTATGAAGGTCAGGCGCCGAACGACCTAAAAATTAA<br>CAAATGGATTGCGCTAGTTCTGTATGCGATTGTAGCGAGTGTTGCGACCTTTGTTCATGGTGGTTTTTCTGGTTGGCA<br>GCCGATTATTTATAAAACAGGTGCGTTTAGCGAACTGTGTAAAGAAGGTGATGAAGTTCAAATTTTTAAAGTTGACGA<br>GAATATTAGCTATAATAGTTGTGGTAATAGAGATGCGGCGATTAATAATCTGTTTACACTGTCGTTCTTTCTGCATTTTT<br>TCCTGAGCAGCATGTCTGGTTTTATTCTGGATACGTTTGGTGAAAAAGTTTGTTTCCTGTGTGGTCAGATTATTCTGGC<br>GCTGGCGTTTTTTAGTCTGGCGATTCTGAAATTCAGCTACGTTTGGTATATTTTCTTTTTTCTGTTAGGTGTTAGCGCG<br>GATTTAAGTTTTATTCCCCTGCTGAAAATTTCGAAATATTTTCCCGGTCAGGAATCGCTGATTTTTGGTATATTAGGTA<br>GCGCGAGAAGTACCGGTTTTGCGGTTGGTCTGCTGTTAAAAATTGCGTTTTTCTATTTATTTAATTTTGGTAATAATGA<br>ATTTTATATTCTGTGTGTTATTTATCTGTGTACATGTATTCTGTTCAGCTTTCTAGTTGGTATTTTTATTATGCCGGGTAA<br>GTCTACCAATAATAAAAGTACCATGCAGGGTCAGGATAATATTCAGATTAATTCTGAACGTGCGACGGGTAGCTCGAT<br>GACAGATAAGAAAAATAATAATAATAATAATAATAATAATAATAATGGTATTTTCCAGGAAATGGAAAATCTGGAATCTG<br>GTAAAAGAAGCAGCATTAATATTAGCGACAGAAATAGCCTGAGTAAACCGCAGAATAATAATAATAATAATACGTCGC<br>TACTGGAATATATAAAAAGTCTGTGGGCGCATCCCCAGAAATGGGAATATCTGATTACAGTTTTTATTTGTTCTAGCTC<br>TATGATTAGATTTGATTATTTTATAAAAACAAATAGAAGTTTCTTTATGACGAATGCGTATGATCTGACAACGGTATTTA<br>GTCTGAGCACAGTTCTAAGTTTTCTGCCCAGTCCCCTATTTGGTTATATTTCTGGTAAACTGGGTAGTGTTTATGGTAT<br>ACTGATTAATAATTTATTTATTCTACTGACATATGTTTGTGTTCTGTTTAATCCGATACTGTTCAAATATATTGCGATATT<br>TACATTTTTCTTCTTCATTAGCTTTGCGTTTAGCTGTTTCTACTGTTATGTTGATGAAAAATATAGCAAGGAACATTTTG<br>GTAAATTATGTGGTATTATGTTTGCGGTTTCTGCGATTTGTTTATCGATAAACTTTTATCTGACATACCTAACAGATGTT<br>GTATATGCGAAACTAGGTGAATTCAAACACATGCCCGTAACGTACGGTTTAAACGTATTAGGTCTAGTTTCGCTGATT<br>GGTTGTATTTATTTAAAGGTTACAGAAAAGAAGAAAAACAAAAACTGA | Codon-harmonized sequence of PfApiAT2 (PF3D7_0914700), for heterologous expression in *Xenopus laevis* oocytes |
| PfApiAT2-mNG_Xl | ATGGCGAGCGATGTTAGCAAGGAAAATATTAGCCAGCTAAAGTATGAAGGTCAGGCGCCGAACGACCTAAAAATTAA<br>CAAATGGATTGCGCTAGTTCTGTATGCGATTGTAGCGAGTGTTGCGACCTTTGTTCATGGTGGTTTTTCTGGTTGGCA<br>GCCGATTATTTATAAAACAGGTGCGTTTAGCGAACTGTGTAAAGAAGGTGATGAAGTTCAAATTTTTAAAGTTGACGA<br>GAATATTAGCTATAATAGTTGTGGTAATAGAGATGCGGCGATTAATAATCTGTTTACACTGTCGTTCTTTCTGCATTTTT<br>TCCTGAGCAGCATGTCTGGTTTTATTCTGGATACGTTTGGTGAAAAAGTTTGTTTCCTGTGTGGTCAGATTATTCTGGC<br>GCTGGCGTTTTTTAGTCTGGCGATTCTGAAATTCAGCTACGTTTGGTATATTTTCTTTTTTCTGTTAGGTGTTAGCGCG<br>GATTTAAGTTTTATTCCCCTGCTGAAAATTTCGAAATATTTTCCCGGTCAGGAATCGCTGATTTTTGGTATATTAGGTA<br>GCGCGAGAAGTACCGGTTTTGCGGTTGGTCTGCTGTTAAAAATTGCGTTTTTCTATTTATTTAATTTTGGTAATAATGA<br>ATTTTATATTCTGTGTGTTATTTATCTGTGTACATGTATTCTGTTCAGCTTTCTAGTTGGTATTTTTATTATGCCGGGTAA<br>GTCTACCAATAATAAAAGTACCATGCAGGGTCAGGATAATATTCAGATTAATTCTGAACGTGCGACGGGTAGCTCGAT<br>GACAGATAAGAAAAATAATAATAATAATAATAATAATAATAATAATGGTATTTTCCAGGAAATGGAAAATCTGGAATCTG<br>GTAAAAGAAGCAGCATTAATATTAGCGACAGAAATAGCCTGAGTAAACCGCAGAATAATAATAATAATAATACGTCGC<br>TACTGGAATATATAAAAAGTCTGTGGGCGCATCCCCAGAAATGGGAATATCTGATTACAGTTTTTATTTGTTCTAGCTC<br>TATGATTAGATTTGATTATTTTATAAAAACAAATAGAAGTTTCTTTATGACGAATGCGTATGATCTGACAACGGTATTTA<br>GTCTGAGCACAGTTCTAAGTTTTCTGCCCAGTCCCCTATTTGGTTATATTTCTGGTAAACTGGGTAGTGTTTATGGTAT<br>ACTGATTAATAATTTATTTATTCTACTGACATATGTTTGTGTTCTGTTTAATCCGATACTGTTCAAATATATTGCGATATT<br>TACATTTTTCTTCTTCATTAGCTTTGCGTTTAGCTGTTTCTACTGTTATGTTGATGAAAAATATAGCAAGGAACATTTTG<br>GTAAATTATGTGGTATTATGTTTGCGGTTTCTGCGATTTGTTTATCGATAAACTTTTATCTGACATACCTAACAGATGTT<br>GTATATGCGAAACTAGGTGAATTCAAACACATGCCCGTAACGTACGGTTTAAACGTATTAGGTCTAGTTTCGCTGATT<br>GGTTGTATTTATTTAAAGGTTACAGAAAAGAAGAAAAACAAAAACATGGTGAGCAAAGGAGAAGAAGATAACATGGCT<br>AGCCTGCCAGCTACACATGAACTGCATATTTTTGGAAGCATTAACGGAGTGGATTTTGATATGGTGGGACAGGGAAC<br>AGGAAACCCAAACGATGGATATGAAGAACTGAACCTGAAAAGCACAAAAGGAGATCTGCAGTTTAGCCCATGGATTC<br>TGGTGCCACATATTGGATATGGATTTCATCAGTATCTGCCATATCCAGATGGAATGAGCCCATTTCAGGCTGCTATGG<br>TGGATGGAAGCGGATATCAGGTGCATAGAACAATGCAGTTTGAAGATGGAGCTAGCCTGACAGTGAACTATAGATAT<br>ACATATGAAGGAAGCCATATTAAAGGAGAAGCTCAGGTGAAAGGAACAGGATTTCCAGCTGATGGACCAGTGATGAC<br>AAACAGCCTGACAGCTGCTGATTGGTGCAGAAGCAAAAAAACATATCCAAACGATAAAACAATTATTAGCACATTTAA<br>ATGGAGCTATACAACAGGAAACGGAAAAAGATATAGAAGCACAGCTAGAACAACATATACATTTGCTAAACCAATGGC<br>TGCTAACTATCTGAAAAACCAGCCAATGTATGTGTTTAGAAAAACAGAACTGAAACATAGCAAAACAGAACTGAACTTT<br>AAAGAATGGCAGAAAGCTTTTACAGATGTGATGGGAATGGATGAACTGTATAAATGA | Codon-harmonized sequence of fusion protein PfApiAT2-mNeonGreen (PF3D7_0914700), for heterologous expression in *Xenopus laevis* oocytes |
