## Supplementary Table 2 for "PfApiAT2 is a proline transporter essential for the transmission of *Plasmodium falciparum* by the mosquito vector"

| Epitope/antigen | Host species | Dilution for IFA | Source |
|---|---|---|---|
| HA | Rabbit | 1 in 200 | RM305, SAB5600116, Sigma-Aldrich |
| HA | Mouse | 1 in 200 | 16B12, Sigma-Aldrich |
| GAPDH | Mouse | 1 in 200 | MABS1946, EMD Millipore |
| Cap380 | Rabbit | 1 in 500 | This study |
| Anti-mouse-568 | Goat | 1 in 1000 | A-11004, Thermo Scientific |
| Anti-mouse-647 | Goat | 1 in 1000 | A-21235, Thermo Scientific |
| Anti-rabbit-546 | Goat | 1 in 1000 | A-11035, Thermo Scientific |
| Anti-rabbit-647 | Goat | 1 in 1000 | A-21244, Thermo Scientific |
