## Supplementary Figure and Table legends for "PfApiAT2 is a proline transporter essential for the transmission of *Plasmodium falciparum* by the mosquito vector"

**Supplementary Figure 1: Parasites recrudescing after HFG-selection with mutations in *pfapiat2* locus show increased survival under HFG pressure.**

Ring stage parasites were assessed for survival under HFG drug pressure in a standard 72hr assay and showed a comparable shift in EC_50_ as previously reported [20] and no shift in survival under control compounds atovaquone (ATQ) and dihydroartemisinin (DHA) (HFG WT vs G449R P<0.0001, WT vs R345I P<0.0001, One-way ANOVA, post-hoc Holm-Šídák's multiple comparisons correction). EC_50_ values from three independent biological replicates, measuring survival over a range of concentrations, with three technical replicates per concentration. All comparisons and details of statistical tests provided in Supplementary Data. Source data are provided as a Source Data file.

**Supplementary Figure 2: HFG-R oocysts do not increase in size over 14 or 21 days of development.**

Oocyst development was assessed in mosquitoes infected with WT, G449R and R345I oocysts at 14 (left) and 21dpi (right). HFG-R oocysts do not show an increase in size, unable to catch up to WT (14dpi WT vs R345I P<0.0001, WT vs G449R P<0.0001; 21dpi 14dpi WT vs R345I P<0.0001, WT vs G449R P<0.0001, Kruskal-Wallis with post-hoc Dunn’s correction). All comparisons and details of statistical tests provided in Supplementary Data. Source data are provided as a Source Data file.

**Supplementary Figure 3: Successful genetic disruption of *pfapiat2* was assessed via PCR around integration site and survival under HFG-pressure.**

A: PCR of integration site, showing location of primers to confirm integration of hDHFR cassette in the *pfapiat2* locus. Gel electrophoresis of PCR products confirmed the presence of transgenic DNA insert in *pfapiat2* locus in the transgenic parasites, but not in the WT, and confirmed the absence of WT DNA in transgenic PfApiAT2-KO parasites. PCR products run on the same gel the same time, cropped for ease of visualizing. B: PfApiAT2-KO asexual blood-stage parasites show increased survival under HFG pressure as previously reported [20] but not under control compounds atovaquone (ATQ) and dihydroartemisinin (DHA) (HFG WT vs PfApiAT2-KO P<0.0001, One-way ANOVA, post-hoc Holm-Šídák's multiple comparisons correction). EC_50_ values from three independent biological replicates, measuring survival over a range of concentrations, with three technical replicates per concentration. All comparisons and details of statistical tests provided in Supplementary Data. Source data are provided as a Source Data file.

**Supplementary Figure 4: Successful integration of smHA tag at the C-terminus of PfApiAT2 and localization to the membrane in asexual blood-stage transgenic parasites.**

A: Schema of integration showing cut site and repair with linearized donor plasmid downstream of the *pfapiat2* locus. B: PCR of integration site, showing location of primers to confirm integration of smHA tag at the C-terminus of the *pfapiat2* locus and the hDHFR downstream. Gel electrophoresis of PCR products confirmed the presence of transgenic DNA insert in the transgenic parasites, but not in the WT, and confirmed the absence of WT DNA in transgenic PfApiAT2-smHA parasites. C: Confocal microscopy IFA of asexual blood-stage PfApiAT2-smHA parasites showing signal when stained with an anti-HA (magenta) antibody at the periphery of the parasite. GAPDH (yellow, cytoplasm), Hoechst (cyan, DNA) and wheat germ agglutinin WGA (white, RBC membrane) also visualized. No HA signal was observed in WT parasites confirming specificity of signal.

**Supplementary Figure 5: Heterologous expression of PfApiAT2 and PfApiAT2-mNG in *X. laevis* oocytes show membrane localization of PfApiAT2-mNG, electro-neutrality during proline flux and a predicted electro-neutral transport pore.**

A: Constructs used for heterologous expression: PfApiAT2 and PfApiAT2-mNeonGreen (mNG) driven by a T7 promoter and stabilized by 5’ and 3’ *Xenopus* globin UTRs upstream and downstream of the gene [29]. B: Membrane localization of PfApiAT2-mNG at the plasma membrane of *Xenopus* oocytes, co-stained with membrane dye FM4-64. C: Proline flux into *Xenopus* oocytes 4 days post-injection with cRNA encoding PfApiAT2 results in no change in membrane potential when probed with either (i) -80mV or (ii) +80mV across the membrane. Data shown is representative of two biological replicates. D: (i) AlphaFold structural prediction of PfApiAT2. (ii, iii) A large electropositive bias is predicted on one face of the protein, likely to be the cytoplasmic side. (iv, v) Sections through the predicted structure (orange overlay in ii and iii) show both electro-positive and electro-neutral transport channel. Source data are provided as a Source Data file.

**Supplementary Figure 6: All PfApiAT2-KO oocysts grow with additional nutrients but are not rescued by supplementation of proline into mosquito sugar meal.**

A: All PfApiAT2-KO oocysts from all mosquitoes show following an additional blood (2BF) or protein meal (BSA) as compared to controls (1BF). We were unable to observe a subset of stalled oocysts that do not respond to additional nutrients. Each dot represents measured area of an individual oocyst, and the same data is depicted as average oocyst size/gut in Figure 6B. B: *Ad libitum* supplementation of proline at 20mM in mosquito sugar meal does not rescue PfApiAT2-KO oocyst size at 7dpi (WT (-) vs PfApiAT2-KO (-) *p*<0.0001; PfApiAT2-KO (-) vs PfApiAT2-KO 20mM Pro *p*>0.9999; WT (-) vs PfApiAT2-KO 20mM Pro *p*>0.0001). Each dot represents average area of oocysts per midgut from midguts with > 2 midguts and line represents median, data assessed using Kruskal-Wallis test with post-hoc Dunn’s correction. Data from two independent biological replicates, all comparisons and details of statistical tests provided in Supplementary Data. Source data are provided as a Source Data file.

**Supplementary Figure 7: The *pfapiat2* locus has fewer nonsynonymous to synonymous mutation rate compared to pan-genome median from clinical isolates.**

A: Nonsynonymous to synonymous mutation values extracted from PlasmoDB were compared for all genes across the parasite genome. Median across the genome (black) is greater than *pfapiat2* (red), confirming purifying selective pressure despite blood-stage dispensability. B: SNPs observed in *pfapiat2* are rare and low abundance. Mutations in *pfapiat2* were extracted from the Pf8 database, and nonsynonymous mutations were found to be very low-abundance in the population indicating a strong conservation of sequence across the locus.

**Supplementary Table 1: Relevant primers and DNA sequences used in this study**

Primers used for cloning, integration PCR and codon-harmonized sequences of PfApiAT2 and PfApiAT2-mNG used for heterologous *Xenopus laevis* oocyte expression.

**Supplementary Table 2: Antibodies used in this study**

All relevant antibodies, their host species, relevant concentration and sources.
